# PyBindingCurve, simulation and curve fitting to complex binding systems at equilibrium

**DOI:** 10.1101/2020.11.06.371344

**Authors:** Steven Shave, Yan-Kai Chen, Nhan T. Pham, Manfred Auer

## Abstract

Understanding multicomponent binding interactions in protein-ligand, protein-protein and competition systems is essential for fundamental biology and drug discovery. Hand deriving equations quickly becomes unfeasible when the number of components is increased, and direct analytical solutions only exist to a certain complexity. To address this problem and allow easy access to simulation, plotting and parameter fitting to complex systems at equilibrium, we present the Python package PyBindingCurve. We apply this software to explore homodimer and heterodimer formation culminating in the discovery that under certain conditions, homodimers are easier to break with an inhibitor than heterodimers and may also be more readily depleted. This is a potentially valuable and overlooked phenomenon of great importance to drug discovery. PyBindingCurve may be expanded to operate on any equilibrium binding system and allows definition of custom systems using a simple syntax. PyBindingCurve is available under the MIT license at: https://github.com/stevenshave/pybindingcurve as Python source code accompanied by examples and as an easily installable package within the Python Package Index.

## Introduction

Simulation and fitting of experimental parameters for systems describing binding at equilibrium is a fundamental need in drug discovery, along with many facets of biology from fundamental to systems. The advantages provided in experimental planning alone justify its importance, allowing expectations of signal strength and species abundance to guide experimental, assay, and instrument setup. In this manuscript, we document the creation of PyBindingCurve and apply it to the most useful protein-ligand systems in biology and drug discovery, deriving equations for the resultant population abundances at equilibrium. Beyond the simplest systems, manual derivation of direct algebraic solutions becomes difficult and we therefore turn to symbolic manipulation in software to derive solutions. These systems are described by polynomial equations with multiple solutions, presenting the problem of not only choosing the correct solution, but choosing the correct solution throughout an experiment where the physically relevant solution may change throughout a titration. In addition, there are no general direct solutions to polynomials of order greater than four (Ayoub, 1980; Rosen, 1995), limiting the scope for deriving direct analytical solutions to highly complex systems. We therefore turn to Lagrange multipliers and root-finding techniques, transforming the problem into one of constrained optimization. We compiled these solutions with methods to simulate titrations, plot results and fit parameters for experimental value determination into a Python package named PyBindingCurve, allowing simple simulation, plotting, and fitting of systems at equilibrium. Multiple freely available and commercial software packages exist for plotting and simulating systems (Munson, 1983; Royer, 1993; Royer et al, 1990a), however most are closed-source, hindering development and integration into existing workflows, or require real solutions to be derived or transferred from literature (Bronstein et al, 1989; Thrall et al, 1996), a difficult process, especially with the added problem of selecting the correct polynomial root representing physically relevant solutions. Having an opensource framework to simulate, fit and interrogate these systems brings with it great advantages allowing automation and integration into existing software pipelines and analysis methods.

All binding events may be described over time by an on-rate (with units: M^-1^·s^-1^), describing the rate of association, and an off-rate describing the complex falling apart into its constituent species (units: s^-1^). The interplay between complex association and dissociation in a closed system creates a dynamic equilibrium containing a steady state of species abundances. We may combine these rate constants describing the population at equilibrium into the dissociation constant; K_D_ (units: M), defined simply as the off-rate divided by the on-rate. Conveniently, when one binding site is present, such as in 1:1 binding, this value denotes the concentration at which 50% of a species will be in complex with its binding partner. Lower values denote higher affinity interactions and therefore tighter binding. Often, we may derive direct analytical solutions to the concentration of complex formed. A full derivation of 1:1 binding from mass balances is available as Supporting Equation 1 and can be found in literature (Green, 1965; Hulme & Trevethick, 2010; Inglese et al, 1989; Wang et al, 1992). Simulation of binding curves with this equation requires no special treatment, with only one polynomial root being physically relevant across all possible experimental parameter values. This provides a direct method for calculating complex concentration over a range of system parameters.

In addition to 1:1 binding, a common system is 1:1:1 competition, commonly used in drug discovery efforts to detect new chemical entities displacing a known binder. As an example, a fluorescently labelled ligand in complex with target protein may have its fluorescence anisotropy measured (Lakowicz, 2006; Weber, 1953). Displacement of the labelled ligand by another inhibitor competing for the same binding site will remove labelled ligand from the complex resulting in a change of anisotropy as a function of complex concentration. The 1:1:1 competition binding equation may be solved in the same manner as demonstrated in supporting material for 1:1 binding with changes made to the mass balances, and results in a third order, or cubic equation requiring significantly more manipulation but remaining feasible by hand. This results in three possible polynomial roots as solutions, one of which may be entirely excluded as never physically relevant, whilst the choice between the other two is dependent on the ligand and inhibitor K_D_s relative to each other. This solution is also readily available in literature (Teukolsky et al, 1992). Additionally, the breaking of heterodimers with an inhibitor can be represented by this 1:1:1 competition system, where protein monomers can be thought of as a protein and ligand which upon binding become a heterodimer whilst the inhibitor competes for a binding site on one of the protein monomers.

Studies in biology often involve oligomers, or repeating units, the conceptually simplest of which is a homodimer; two identical monomer units binding to each other. Homodimers have been characterized as having distinct properties setting them apart from heterodimers which comprise two chemically different proteins. In general, the binding interfaces of homodimers are larger with more interacting residues, specifically enriched with hydrophobic residues (Zhanhua et al, 2005). Mathematically understanding this construct and its behavior is critically important for fundamental biology. It is therefore surprising that we were able to find only one instance in literature of derivation of a direct, analytical solution to homodimer formation at equilibrium (Benfield et al, 2011). Upon first considering homodimer formation, it appears a simpler case than 1:1 protein ligand binding as only one starting species is involved, two units of which transition to become a single dimer upon complexation. An important consideration exists when the dimer undergoes dissociation and two monomers are produced, increasing the concentration of free monomer at double the dissociation rate. The same is true in reverse for complexation; with two monomers consumed for creation of one dimer. The derivation of a direct analytical solution to dimer formation can be found in the supporting information accompanying this manuscript and is labelled Supporting Equation 2. Like the solution of the quadratic for 1:1 binding, one polynomial root is always physically correct.

We were unable to find literature providing direct analytical solutions to equations describing homodimer breaking with an inhibitor. This is surprising as fundamental biology and drug discovery efforts often seek to deactivate or break apart homodimers with small molecules or peptides mimicking interaction surfaces. As the ligand and protein are the same species, manual derivation of direct analytical solutions for amount dimer formed proves difficult. At this level of complexity, direct analytical solutions can be found using symbolic manipulation with programs such as Wolfram Mathematica (Wolfram Research, 2019). See supporting information codes 1, 2, 3, and 4 for Wolfram Mathematica code solving the 1:1, 1:1:1 competition, dimer formation and dimer breaking systems respectively. In highly complex systems, such as homodimer breaking, we may observe the physically correct solution “switching” from one polynomial root to another as titrations progress. We have termed this ‘solution switching’, requiring tracing steps to be built into PyBindingCurve allowing smooth transition from point to point when using the derived direct analytical solutions to complex systems. To simulate and perform fitting such as K_D_ determination using experimental data using systems which for which no direct analytical solutions exist, we use Lagrange multipliers in an approach similar to that taken by Royer (Royer et al, 1990b) to solve the systems expressed as constrained optimization problems (See Supporting Code Listing 5 to 8). PyBindingCurve allows custom systems to be defined and solved using Lagrangian based techniques, simply specifying “P+P<->PP” defines homodimer formation, whilst “P+L<->PL, P+I<->PI” defines 1:1:1 competition (see “Simulation of custom binding systems” in the accompanying supporting material). Additionally, systems may be solved kinetically as a system of ordinary differential equations (ODEs - see Supporting Code Listings 9-12). PyBindingCurve automatically utilizes the fast, direct analytical solutions to systems where possible, otherwise the Lagrangian approach to constrained optimization is used. Kinetic solvers are not used by default but can be specified. Bellow, we document the result of using PyBindingCurve to explore the striking differences between homo- and hetero-dimer breaking and its implications. The PyBindingCurve software package available at https://github.com/stevenshave/pybindingcurve allows simulation, fitting and derivation of systems parameters for a range of predefined and custom definable systems.

## Results and discussion

Using the PyBindingCurve package we investigated the large and surprising differences between homo- and hetero-dimer formation. Understanding of these differences has the potential to impact fundamental biology, with better use of tool compounds and an improved systems biology-based understanding of biological pathways. We must first consider dimer formation, best illustrated by a theoretical experiment where an increasing concentration of monomer is titrated. Whilst experimentally it is not easy to increase monomer concentration, the experiment could be performed in reverse with buffer titrated into a known starting amount of monomer and dimer concentration monitored. Dimer half-life would need to be considered, ensuring an equilibrium is achieved before each dimer measurement is recorded. Performing the experiment for heterodimers is the simplest conceptually, whereby a known, and equal starting concentration of each monomer ‘A’ and ‘B’ is diluted, giving a total number of particles in a constant volume of N. To directly compare homodimer complexation with this system, we must use twice the homodimer ‘H’ monomer concentration to achieve the same number of particles as N.

**Error! Reference source not found**.**Fig 1**A, shows dimer formation as a function of monomer for dimers with a dissociation constant (K_D_) of 100 nM. It is evident that with the same number of particles present, more homodimer than heterodimer is formed. This is expected as homodimer monomers, can form a dimer with any other monomer. Heterodimers are only formed when two complimentary monomers come together, effectively halving the concentration of binding partners that a heterodimer monomer encounters. Fig 1B illustrates profound differences observed upon dimer breaking with an inhibitor, the understanding of which is of crucial importance for drug discovery and fundamental biology involving the application of drug and tool compounds for dimer breaking. We can observe a system where dimer complexation starting with a total of 2 µM homodimer monomer with a dimerization K_D_ of 100 nM has an inhibitor titrated into it with a K_D_ of 10 nM to homodimer monomer. Comparing this to a similar system containing 1 µM of both components A and B of a heterodimer monomer (total 2 µM monomer concentration) with a similar dimerization inhibitor with a K_D_ of 10 nM to A, we observe striking differences. First as expected from the dimer formation plot in Figure 1A, the starting dimer concentration of homodimer is higher than that of heterodimer. Again, this can be explained simply by the fact that a homodimer monomer can bind any other homodimer monomer, whereas with a heterodimer monomer, a monomer must encounter a complimentary monomer for complexation to occur. As the titration continues, an interesting crossover occurs at around 4 µM inhibitor concentration with an equal amount of homo- and heterodimer present. As inhibitor concentration increases, the amount of heterodimer present is greater than that of homodimer. This effect can be explained by the increased abundance of inhibitor binding partner in the case of homodimers. As in the previous explanation, particles of homodimer are encountered at twice the rate of particles of appropriate heterodimer monomer by the inhibitor diffusing in solution, causing more homodimer monomer-inhibitor complex than heterodimer monomer-inhibitor complex. For the same amount of inhibitor in solution, a greater proportion complexes with monomer in the case of homodimers, removing free monomer capable of complexing with another monomer into a dimer. This is interesting, as initially, the increased dimerization of homodimer is greater than the effect of increased homodimer monomer-inhibitor formation. There is however, a point in inhibitor titration where this balance shifts, and the effect of the increased homodimer monomer-inhibitor formation is greater than the increased dimerization, shifting the dynamic such that homodimers may be more easily broken apart by inhibitors. A final observation can be made, in that the depletion of homodimer is more complete than that of heterodimers at high concentrations of inhibitor. Again, this can be explained by the abundance of inhibitor binding partners, each monomer in the case of homodimers, versus one monomer in the case of heterodimers. Further exploring the switch of homo- versus hetero-dimer ease of breaking, we may visualize areas of affinity space where breaking one is easier than the other, as shown in Fig 2.

**Fig 1.**
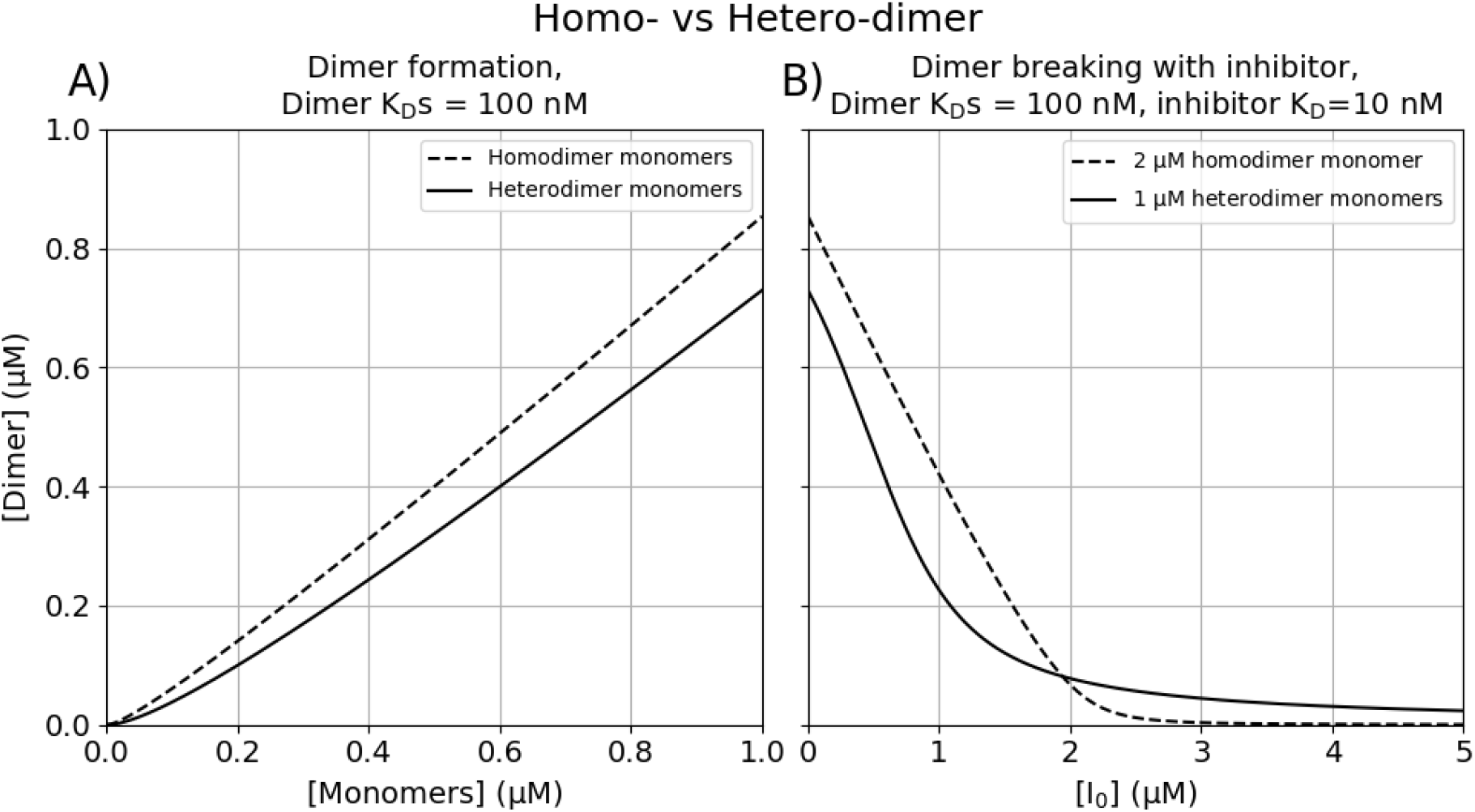
Homo-vs-Heterodimer making and breaking. A) Dimer formation with a K_D_ of 100 nM as a function of monomer concentration. As a homodimer (broken line) contains two copies of the same monomer, total monomer concentration is twice that shown on the x-axis. Heterodimer formation (solid line) contains two different monomers with concentration for each monomer given by the x-axis. B) After monomer titration to 1 µM as shown in A, or effectively 2 µM in the homodimer case, the resultant complex has an inhibitor (I_0_) titrated against it. The inhibitor has a K_D_ of 10 nM to one heterodimer monomer (solid line), and the same K_D_ to homodimer monomers.

**Fig 2.**
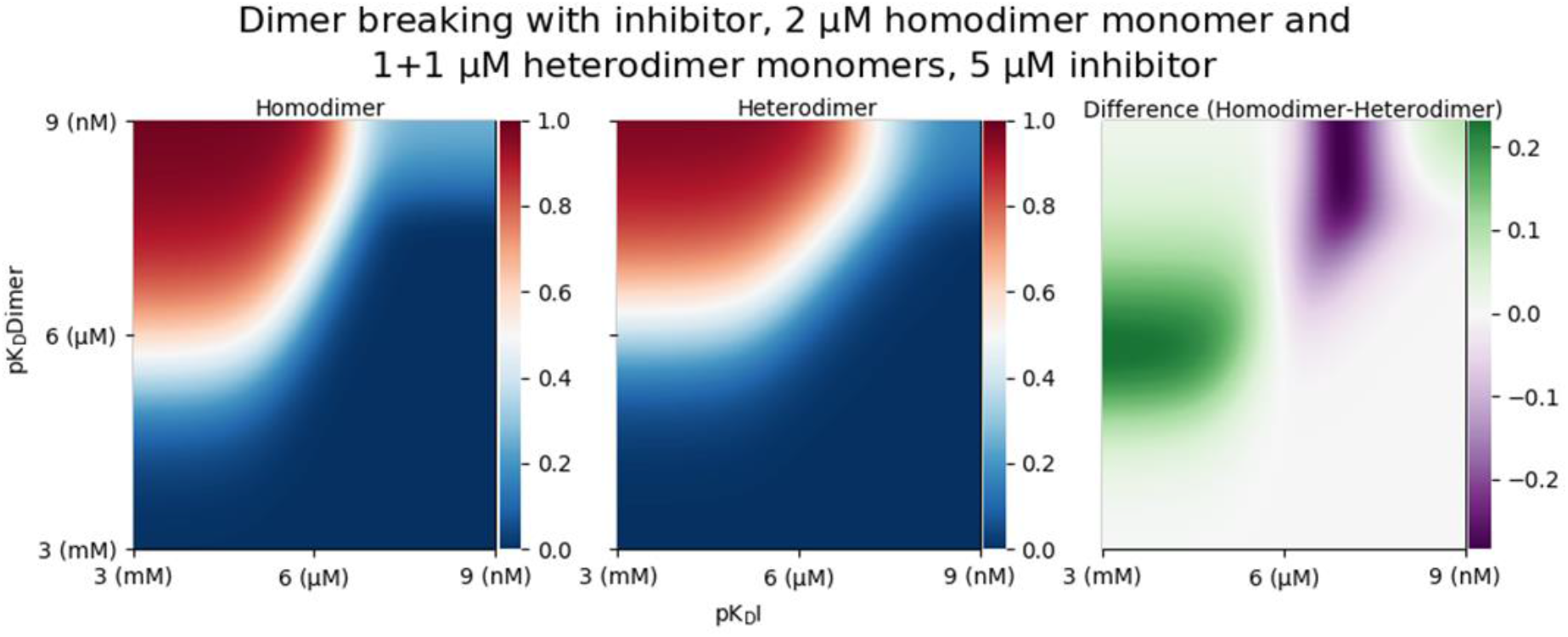
Homodimer versus heterodimer breaking heatmaps showing dimer concentration as a function of changing dimer and inhibitor affinity.

Fig 2 left shows concentration of homodimer formed with a starting concentration of 2 µM monomer and a range of inhibitor and dimerization K_D_s along the x- and y-axes respectively, expressed as pK_D_s. Fig 2 centre shows the same for heterodimer, with 1+1 µMs of the two monomers as a starting concentration and a range of inhibitor K_D_s. Fig 2 right shows the difference between homodimer and heterodimer, green indicating that the homodimer was harder to break, and magenta indicating that it was easier to break.

## Discussion

In development of PyBindingCurve, we iterated through many approaches to simulating protein-ligand systems as their complexity increased. Starting with hand crafted direct analytical solutions, increased system complexity led to computer-generated code requiring tracing approaches to choose the correct solution. As complexity increased and solutions to high order polynomials were no longer found (Ayoub, 1980; Rosen, 1995), such as in 1:4 protein-ligand binding, we transitioned to first using iterative kinetic models describing systems as ODEs, to Lagrange multiplier minimization problems expressed as constrained system optimization problems. PyBindingCurve automatically chooses the most appropriate method to solve common binding systems. The use of Lagrange multipliers led to methods capable of solving user defined binding systems specified in simple text, allowing applicability of PyBindingCurve to practically all biological systems. Having such capabilities present in an open source package promotes freedom to use, integrate and improve PyBindingCurve.

The striking difference in behavior of homodimers versus heterodimers could have a significant impact on drug discovery efforts. Simply dissemination of the knowledge that near complete depletion of homodimers is easier than with heterodimers is valuable, before making any numerical predictions or analysis. We believe a major strength of PyBindingCurve is direct programmatic access for exploration of these binding systems in Python, currently one of the most popular and fastest growing programming languages. This helps allow insights as demonstrated in the homo- versus hetero-dimer formation example which would not have been easily discovered using existing offerings of traditional curve fitting and simulation software.

We envision the continued growth and development of PyBindingCurve. A detailed user guide with tutorials and API documentation is available in supporting information accompanying this manuscript and online (https://stevenshave.github.io/pybindingcurve/).

## Materials and Methods

We implemented PyBindingCurve in Python (version 3.6.8), developing methods capable of simulating, plotting and fitting parameters to experimental results for 1:1, 1:n (where n is 1-5), 1:1:1 competition, homodimer and heterodimer systems. In the testing and validation process for these analytically solved complex systems solved with direct methods, we encountered considerable numerical instability (Goldberg, 1991) and so, use arbitrary precision arithmetic to a high degree of accuracy using the mpmath (version 1.1.0) package (Johansson, 2018). Internally, PyBindingCurve uses a combination of direct analytical solutions for simple binding systems (computationally quick to simulate), and Lagrangian multiplier-based approach to solving constrained systems when dealing with more complex systems. To derive the Python code for simple systems with direct analytical solutions, we used Wolfram Mathematica to derive solutions from sets of mass balances and binding equations (See Supporting Code Listings 1-4). These solutions were written out and the program MathematicaEquationToPython used to convert the equations to Python functions (available at: https://github.com/stevenshave/MathematicaEquationToPython). Systems which proved unsolvable using symbolic manipulation in Mathematica are integrated into PyBindingCurve using Lagrangian multiplier-solved constrained optimisation. Supporting Code Listings 5-8 illustrates the Python code used to construct these Lagrangian systems and simulate 1:1, 1:1:1 competition, homodimer formation, and homodimer breaking. Code present within PyBindingCurve is also capable of parsing custom defined systems and transforming them into Lagrangian functions which are easily solvable. These systems are solved using the fsolve method from the SciPy optimize package. In addition to system simulation at a single set of starting conditions, we created helper functions to enable plotting over a range of species concentrations. Parameter fitting is achieved using the LMFit package (Newville et al, 2016) enabling the calculation of parameters such as K_D_ from experimental data. Full source code along with examples for every system are available in the public GitHub repository accessible at https://github.com/stevenshave/pybindingcurve. Additionally, the PyBindingCurve package has also been submitted to the Python Package Index (https://pypi.org/project/pybindingcurve/) may be installed via pip using the command “pip install pybindingcurve”. Online documentation along with a tutorial is also available at https://stevenshave.github.io/pybindingcurve/.

## Acknowledgements

This Manuscript is dedicated to the memory of the late Dr. Kurt Muller, whose initial work into the automated derivation of systems equations from mass balances and binding equations made this work possible.

The authors acknowledge financial support from the Scottish Universities Life Sciences Alliance (SULSA-http://www.sulsa.ac.uk) and the Medical Research Council (MRC-www.mrc.ac.uk, J54359) Strategic Grant. M.A and N.T.P acknowledge financial support from the Wellcome Trust (Grant 201531/Z/16/Z).

## Author contributions

SS programmed PyBindingCurve with contributions from YKC. SS and NTP contributed to fitting techniques and management of solution switching. MA and SS conceived the project.

