## Supporting Information for "PyBindingCurve, simulation and curve fitting to complex binding systems at equilibrium"

PyBindingCurve source code available at https://github.com/stevenshave/pybindingcurve

### Supporting Equation 1; deriving the fraction ligand bound for 1:1 binding systems.

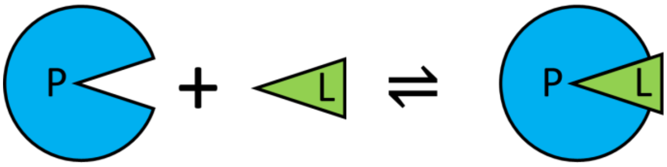

$$[P] = [P_f] + [PL]$$

$$[L] = [L_f] + [PL]$$

$$K_D = \frac{[P_f].[L_f]}{[PL]}$$

$$K_D = \frac{([P] - [PL]).([L] - [PL])}{[PL]}$$

$$K_D[PL] = ([P] - [PL])([L] - [PL])$$

$$K_D[PL] = [P][L] - [P][PL] - [L][PL] + [PL]^2$$

$$0 = [P][L] - K_D[PL] - [P][PL] - [L][PL] + [PL]^2$$

$$0 = [P][L] - [PL](K_D + [P] + [L]) + [PL]^2$$

$$[PL]^2 - ([P] + [L] + K_D)[PL] + [P][L] = 0$$

The quadratic equation above can be solved using the quadratic formula and the helper variables:

$$a = 1,$$

$$b = -(P + L + K_D),$$

$$c = [P][L]$$

Resulting in

$$[PL] = \frac{([P] + [L] + K_D) \pm \sqrt{([P] + [L] + K_D)^2 - 4[P][L]}}{2}$$

**Note: the negative root of the above equation is the physically relevant solution.**

Where [P] is the total protein concentration, [$P_f$] is the free (unbound) protein concentration, [L] is the total ligand concentration, [$L_f$] is the free (unbound) ligand concentration, [$K_D$] is the protein-ligand complex dissociation constant ($K_D$), and [PL] is the protein-ligand complex concentration.

Supporting Equation 2; deriving the fraction total possible dimer present in the dimer formation system

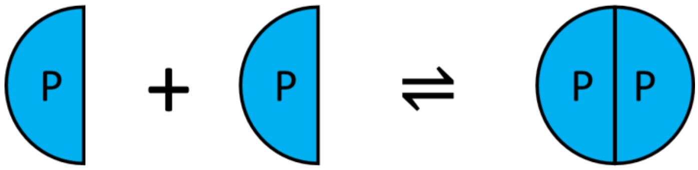

$$[P] = [P_f] + 2[PP]$$

$$K_D = \frac{[P_f].[P_f]}{[PP]}$$

$$K_D = \frac{([P] - 2[PP])([P] - 2[PP])}{[PP]}$$

$$K_D[PP] = ([P] - 2[PP])([P] - 2[PP])$$

$$K_D[PP] = [P]^2 - 4[P][PP] + 4[PP]^2$$

$$4[PP]^2 - 4[P][PP] + [P]^2 - K_D[PP] = 0$$

$$4[PP]^2 - (4[P] + K_D)[PP] + [P]^2 = 0$$

The quadratic equation above can be solved using the quadratic formula and the helper variables:

$$a = 4,$$

$$b = -(4[P] + K_D),$$

$$c = [P]^2$$

Resulting in

$$[PP] = \frac{4[P] + K_D \pm \sqrt{(4[P] + K_D)^2 - 16[P]^2}}{8}$$

**Note: similar to 1:1 binding, the negative root of the above equation is the physically relevant solution.**

Where [P] is the total protein (monomer) concentration, [Pf] is the free (unbound) protein concentration, [$K_D$] is the dissociation constant ($K_D$) of the dimer, and [PP] is the dimer complex concentration.

A similar derivation is also available in Supporting material of: Benfield CT, Mansur DS, McCoy LE, Ferguson BJ, Bahar MW, Oldring AP, Grimes JM, Stuart DI, Graham SC, Smith GL (2011) Mapping the I κ B kinase β (IKKβ)-binding interface of the B14 protein, a vaccinia virus inhibitor of IKKβ-mediated activation of nuclear factor κ B. J Biol Chem 286: 20727-20735

### Supporting Code Listing 1; Mathematica code solving fraction ligand bound to protein in a 1:1 binding system

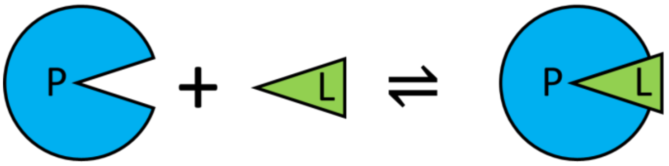

```
Clear["Global`*"]
eqtn = {
  p == pf + pl,
  l == lf + pl,
  pl*Kdpl == pf*lf,
   y == ymin + ((ymax - ymin)*pl)/l
  };
toEliminate = {pf, lf, pl};
Solve[Simplify[Eliminate[eqtn, toEliminate]], y]
```

| Symbol | Meaning |
|---|---|
| `p` | Total protein concentration |
| `pf` | Free protein concentration |
| `l` | Total ligand concentration |
| `lf` | Free ligand concentration |
| `pl` | Protein-ligand complex concentration |
| `Kdpl` | The $K_D$ for the protein-ligand complex |
| `ymin` | The minimum baseline readout signal |
| `ymax` | The maximum readout intensity |
| `y` | Fraction ligand bound to protein |

Output:

$$\Big\{\Big\{y \to -\frac{1}{2\,l}\Big(-(\mathrm{ymax}-\mathrm{ymin})\sqrt{\mathrm{Kdpl}^2+2\,\mathrm{Kdpl}\,l+2\,\mathrm{Kdpl}\,p+l^2-2\,l\,p+p^2}- \mathrm{Kdpl}\,\mathrm{ymax}+\mathrm{Kdpl}\,\mathrm{ymin}-l\,\mathrm{ymax}-l\,\mathrm{ymin}-p\,\mathrm{ymax}+p\,\mathrm{ymin}\Big)\Big\},$$

$$\Big\{y \to -\frac{1}{2\,l}\Big((\mathrm{ymax}-\mathrm{ymin})\sqrt{\mathrm{Kdpl}^2+2\,\mathrm{Kdpl}\,l+2\,\mathrm{Kdpl}\,p+l^2-2\,l\,p+p^2}-\mathrm{Kdpl}\,\mathrm{ymax}+ \mathrm{Kdpl}\,\mathrm{ymin}-l\,\mathrm{ymax}-l\,\mathrm{ymin}-p\,\mathrm{ymax}+p\,\mathrm{ymin}\Big)\Big\}\Big\}$$

### Supporting Code Listing 2; Mathematica code solving fraction ligand bound to protein in a 1:1:1 competition binding system

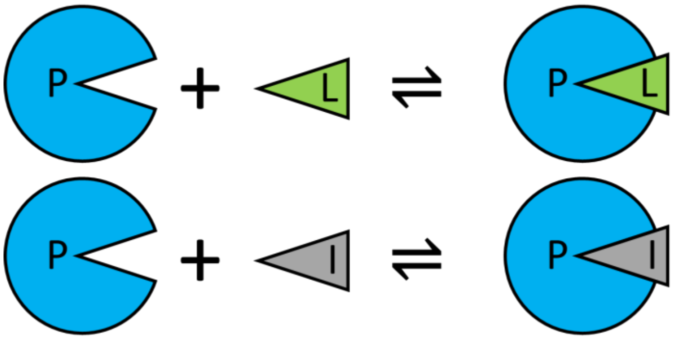

```
Clear["Global`*"]
eqtn = {
   p == pf + pl + pi,
   i == if + pi,
   l == lf + pl,
   pl*Kdpl == pf*lf,
   pi*Kdpi == pf*if,
   y == ymin + ((ymax - ymin)*pl/l)};
toEliminate = {pf, lf, if, pi, pl};
Solve[Simplify[Eliminate[eqtn, toEliminate]], y]
```

| Symbol | Meaning |
|---|---|
| p | Total protein concentration |
| pf | Free protein concentration |
| i | Total inhibitor concentration |
| if | Free inhibitor concentration |
| l | Total (labelled) ligand concentration |
| lf | Free (labelled) ligand concentration |
| pl | Protein-ligand complex concentration |
| pi | Protein-inhibitor complex concentration |
| Kdpl | The $K_D$ for the protein-ligand complex |
| Kdpi | The $K_D$ for the protein-inhibitor complex |
| ymin | The minimum baseline readout signal |
| ymax | The maximum readout intensity |
| y | Fraction ligand bound to protein |

Output too large to list here, can be found converted to Python code in the source code at:

https://github.com/stevenshave/pybindingcurve/blob/master/pybindingcurve/systems/analytical_equations.py

### Supporting Code Listing 3; Mathematica solving the fraction total possible dimer present in the dimer formation system

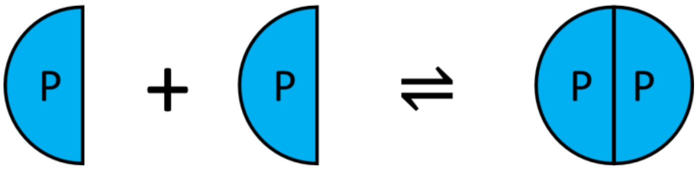

```
Clear["Global`*"]
eqtn = {
   p == pf + (2*pp),
   pp*Kdpp == pf*pf,
   y == ymin + ((ymax - ymin)*(2*pp/p))};
toEliminate = {pf, pp};
Solve[Simplify[Eliminate[eqtn, toEliminate]], y]
```

| Symbol | Meaning |
|---|---|
| p | Total protein concentration |
| pf | Free protein concentration |
| pp | Total dimer concentration |
| Kdpp | The $K_D$ for the protein dimer complex |
| ymin | The minimum baseline readout signal |
| ymax | The maximum readout intensity |
| y | Fraction possible dimer |

Output:

$$\left\{\left\{y \to \frac{1}{4}\left(-\frac{\sqrt{\text{Kdpp}}\ \sqrt{\text{Kdpp} + 8\,p\,(\text{ymax} - \text{ymin})}}{p} + \frac{\text{Kdpp ymax}}{p} - \frac{\text{Kdpp ymin}}{p} + 4\,\text{ymax}\right)\right\},\right.$$

$$\left.\left\{y \to \frac{1}{4}\left(\frac{\sqrt{\text{Kdpp}}\ \sqrt{\text{Kdpp} + 8\,p\,(\text{ymax} - \text{ymin})}}{p} + \frac{\text{Kdpp ymax}}{p} - \frac{\text{Kdpp ymin}}{p} + 4\,\text{ymax}\right)\right\}\right\}$$

Supporting Code Listing 4; Mathematica solving the fraction total possible dimer present in the dimer formation system in the presence of a dimerization inhibitor

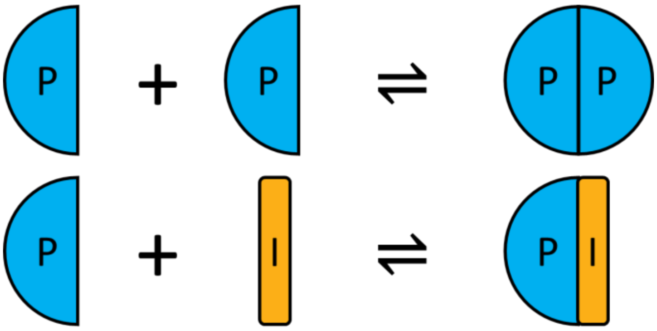

```
Clear["Global`*"]
eqtn4 = {
   p == pf + (2*pp) + pi,
   i == if + pi,
   pp*Kdpp == pf*pf,
   pi*Kdpi == pf*if,
   y == ymin + ((ymax - ymin)*(2*pp/p))};
toEliminate = {pf, if, pi, pp};
Solve[Simplify[Eliminate[eqtn4, toEliminate]], y]
```

| Symbol | Meaning |
|---|---|
| p | Total protein concentration |
| pf | Free protein concentration |
| pp | Total dimer concentration |
| i | Total inhbitor concentration |
| if | Free inhibitor concentration |
| pi | Protein-inhibitor complex concentration |
| Kdpp | The $K_D$ for the protein dimer complex |
| Kdpi | The $K_D$ for the protein-inhibitor complex |
| ymin | The minimum baseline readout signal |
| ymax | The maximum readout intensity |
| y | Fraction ligand bound to protein |

Output too large to list here, can be found converted to Python code in the source code at:

https://github.com/stevenshave/pybindingcurve/blob/master/pybindingcurve/systems/analytical_equations.py

Supporting Code Listing 5; Python code solving 1:1 binding at equilibrium using Lagrangian multiplier-based optimisation of a constrained system and returning protein-ligand complex concentration.

```
from scipy.optimize import fsolve
from autograd import grad
def lagrange_one_to_one(p0,l0,kd):
    def F(X): # Augmented Lagrange function
        response=(X[0]*X[1])/kd
        return response - X[2] *(p0-(response+X[0])) - X[3]*(l0-(response+X[1]))
    dfdL = grad(F) # Gradients of the Lagrange function
    print(dir(dfdL.__str__))
    print(dfdL.__str__())
    pf, lf, lam1, lam2 = fsolve(dfdL, [p0, l0, 1.0,1.0])
    return (pf*lf)/kd
```

Supporting Code Listing 6; Python code solving 1:1:1 binding at equilibrium using Lagrangian multiplier-based optimisation of a constrained system and returning protein-ligand complex concentration.

```
from scipy.optimize import fsolve
from autograd import grad
def lagrange_competition(p0,l0, i0,kdpl, kdpi):
    def F(X): # Augmented Lagrange function
        response=(X[0]*X[1])/kdpl
        constraint1=p0-(response+X[0]+(X[0]*X[2]/kdpi))
        constraint2=l0-(response+X[1])
        constraint3=i0-(X[2]+(X[0]*X[2]/kdpi))
        return response - X[3] *constraint1 - X[4]*constraint2 -X[5]*constraint3
    dfdL = grad(F, 0) # Gradients of the Lagrange function
    pf, lf, inhf, lam1, lam2,lam3 = fsolve(dfdL, [p0, l0, i0, 1.0,1.0,1.0])
    return (pf*lf)/kdpl
```

Supporting Code Listing 7; Python code solving homodimer complexation at equilibrium using Lagrangian multiplier-based optimisation of a constrained system and returning dimer complex concentration.

```
from scipy.optimize import fsolve
from autograd import grad
def lagrange_homodimerformation(p0,kdpp):
    def F(X): # Augmented Lagrange function
        response=((X[0]*X[0])/kdpp)
        constraint1=p0-(response*2+X[0])
        return response - X[1] *constraint1
    dfdL = grad(F, 0) # Gradients of the Lagrange function
    pf, lam1 = fsolve(dfdL, [p0, 1.0])
    return pf*pf/kdpp
```

Supporting Code Listing 8; Python code solving homodimer breaking with an inhibitor using Lagrangian multiplier-based optimisation of a constrained system and returning dimer complex concentration.

```
from scipy.optimize import fsolve
from autograd import grad
def lagrange_homodimerbreaking(p0,i0,kdpp, kdpi):
    def F(X): # Augmented Lagrange function
        pp=((X[0]*X[0])/kdpp)
        pi=((X[0]*X[1])/kdpi)
        constraint1=p0-(pp*2+pi+X[0])
        constraint2=i0-(pi+X[1])
        return pp - X[2] *constraint1 - X[3]*constraint2
    dfdL = grad(F, 0) # Gradients of the Lagrange function
    pf, i_f, lam1,lam2 = fsolve(dfdL, [p0, i0, 1.0,1.0])
    return pf*pf/kdpp
```

Supporting Code Listing 9; Python code solving 1:1 binding at equilibrium kinetically and returning protein-ligand complex concentration.

```
import numpy as np
from scipy.integrate import solve_ivp
def system01_p_l_Kd__pl(parameters: dict, interval=(0, 100)):
    p, l, Kdpl = np.float64(parameters['p']), np.float64(parameters['l']),
np.float64(parameters['Kdpl'])
    def ode(concs, t, Kdpl):
        p, l, pl = concs
        r1 = -p*l + Kdpl*pl
        dpdt = r1
        dldt = r1
        dpldt = - r1
        return [dpdt, dldt, dpldt]
    return solve_ivp(lambda t,y:ode(y,t,Kdpl), interval, [p,l,0.0]).y[2,-1]
```

Supporting Code Listing 10; Python code solving 1:1:1 binding at equilibrium kinetically and returning protein-ligand complex concentration.

```
import numpy as np
from scipy.integrate import solve_ivp
def system02_p_l_i_Kdpl_Kdpi__pl(parameters: dict, interval=(0, 100)):
    p, l, i, Kdpl, Kdpi = np.float64(parameters['p']), np.float64(parameters['l']),
np.float64(parameters['i']), np.float64(parameters['Kdpl']),
np.float64(parameters['Kdpi'])
    def ode(concs, t, Kdpl, Kdpi):
        p, l, i, pl, pi  = concs
        r1 = -p*l + Kdpl*pl
        r2 = -p*i + Kdpi*pi
        dpdt = r1 + r2
        dldt = r1
        didt = r2
        dpldt = - r1
        dpidt = - r2
        return [dpdt, dldt, didt, dpldt, dpidt]
```

Supporting Code Listing 11; Python code solving homodimer complexation at equilibrium kinetically and returning dimer complex concentration.

```
import numpy as np
from scipy.integrate import solve_ivp
def system03_p_Kdpp__pp(parameters: dict, interval=(0, 100)):
    p, Kdpp = np.float64(parameters['p']), np.float64(parameters['Kdpp'])
    def ode(concs, t, Kdpp):
        p, pp  = concs
        r1 = -(p*p) + Kdpp*pp
        dpdt = 2*r1
        dppdt = - r1
```

```
        return [dpdt, dppdt]
    return solve_ivp(lambda t,y:ode(y,t,Kdpp), interval, [p,0.0]).y[1,-1]
```

### Supporting Code Listing 12; Python code solving homodimer breaking with an inhibitor kinetically and returning dimer complex concentration.

```
import numpy as np
from scipy.integrate import solve_ivp
def system04_p_i_Kdpp_Kdpi__pp(parameters: dict, interval=(0, 100)):
    p, i, Kdpp, Kdpi = np.float64(parameters['p']), np.float64(parameters['i']),
np.float64(parameters['Kdpp']), np.float64(parameters['Kdpi'])
    def ode(concs, t, Kdpp, Kdpi):
        p, i, pp, pi  = concs
        r_pp = -(p*p) + Kdpp*pp
        r_pi = -p*i + Kdpi*pi
        dpdt = 2*r_pp + r_pi
        didt = r_pi
        dppdt = -r_pp
        dpidt = -r_pi
        return [dpdt, didt, dppdt, dpidt]
    return solve_ivp(lambda t,y:ode(y,t,Kdpp, Kdpi), interval, [p, i, 0.0, 0.0]).y[2,-1]
```

### PyBindingCurve User guide

PyBindingCurve is a Python package for simulation, plotting and fitting of experimental parameters to protein-ligand binding systems at equilibrium.  In simple terms, the most basic functionality allows simulation of a two species binding to each other as a function of their concentrations and the dissociation constant ($K_D$) between the two species.

#### Installation

PyBindingCurve may be installed from source present in the GitHub repository https://github.com/stevenshave/pybindingcurve via git pull, or from the Python Package Index (https://pypi.org/project/pybindingcurve/) using the command :

```
pip install pybindingcurve
```

#### Requirements

PyBindingCurve was developed using python 3.7.1 but should work with any Python version 3.6 or greater. The following packages are also required

- matplotlib (2.x)
- numpy (1.15.x)
- lm_fit (1.0.0)
- mpmath (1.1.0)
- autograd (1.3)

#### Licence

MIT License

#### Tutorial

##### Simulation of 1:1 binding and exploration of all options

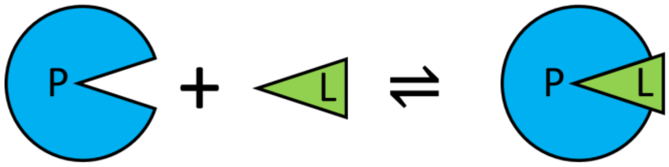

Example code is available here:
https://github.com/stevenshave/pybindingcurve/blob/master/example_1to1_simulation.py

A 1:1 binding system typically consists of a protein and a ligand. However, this 1:1 system is suitable for simulation of any two different species forming a complex. This system can therefore be used to simulate hetero-dimer formation, where two different proteins form a complex. In this simple example, we will imagine wanting to know the concentration of a complex formed. We can choose to work in a common unit, typically nM, or µM, as long as all numbers are in the same unit, the result is valid.  We assume µM for all concentrations and $K_D$s bellow.

First, we may want to produce a plot of protein vs complex concentration for a fixed amount of ligand, simulating the titration of protein into a cuvette for example. We must import PyBindingCurve and NumPy, and then make a new BindingCurve object which takes as an argument the type of system that should be represented. In this case "1:1" will define the correct system. For a list of all systems, please see "pbc.systems and shortcut strings" in the PyBindingCurve documentation.

```
import numpy as np
import pybindingcurve as pbc
my_system = pbc.BindingCurve("1:1")
```

We then define the system parameters in a python dictionary, defining p (protein concentration) as a linear NumPy sequence from 0 to 20 µM, l (ligand concentration) as 10 µM, and the protein-ligand $K_D$ to be 1 µM.

```
system_parameters = {"p": np.linspace(0, 20), "l": 10, "kdpl": 1}
```

We can now add the curve to the plot. If we want multiple simulations on the same plot, then it is good to give the curve a name with the optional name parameter.

```
my_system.add_curve(system_parameters, name= "Curve 1")
```

Now we add another, higher affinity curve to the plot, defining a similar system with a lower $K_D$ and add it to the curve.

```
system_parameters_higher_affinity = {"p": np.linspace(0, 20), "l": 10, "kdpl": 0.5}
my_system.add_curve(system_parameters_higher_affinity, name "Curve 2")
```

Finally show the plot.  Optionally, title, xlabel and ylabel variables may also be passed to title the plot and axes.

```
my_system.show_plot()
```

This produces the following plot:

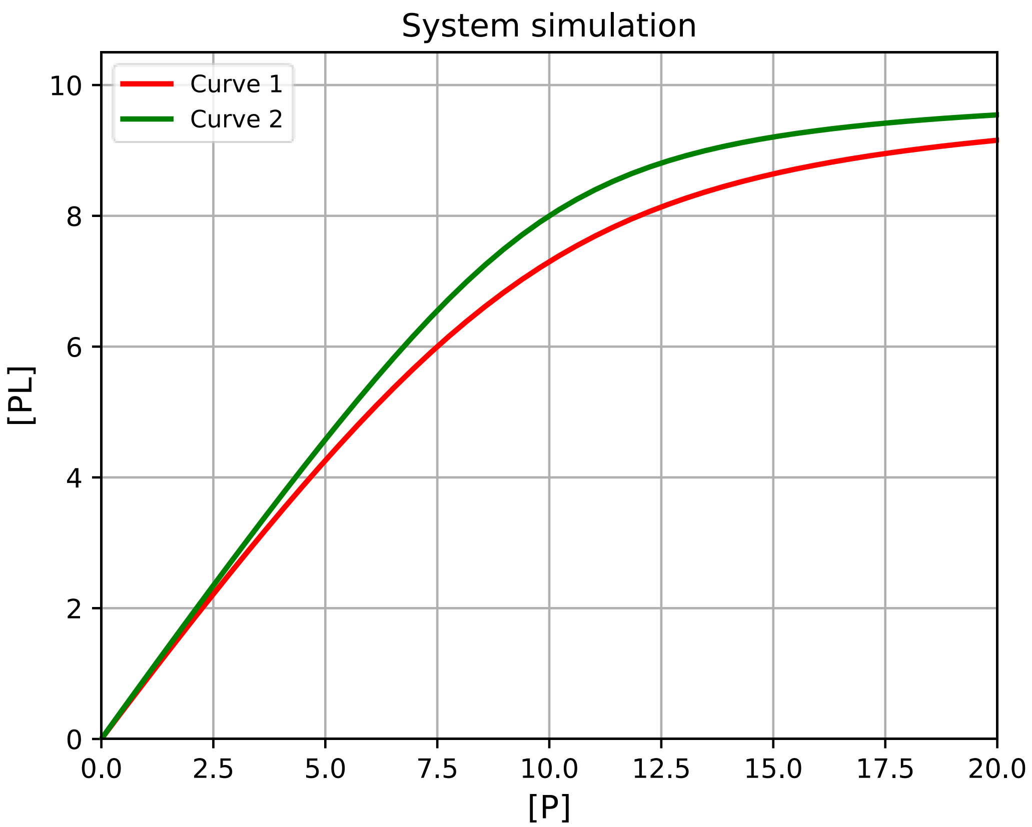

To obtain exact single points from the plot, we may call the query function of my_system:

```
print(my_system.query({“p”:5, “l”:10, “kdpl”:1})
```

If a list or NumPy array is included as a system parameter, then a NumPy array of results is returned.

We may want to simulate a system in terms of a signal, not the concentration of complex. In this case, we may pass additional parameters, setting the ymax and/or ymin variables in the system parameters. Inclusion of these will scale the signal present between these values. This is very important if a detector is used with a maximum or minimum sensitivity and you wish to simulate response. Changing our system parameters to include a maximal response of 2000 units will result in subtle scale and axes changes:

```
system_parameters = {"p": np.linspace(0, 20), "l": 10, "kdpl": 1, 'ymax':2000}
system_parameters_higher_affinity = {"p": np.linspace(0, 20), "l": 10, "kdpl": 0.5,
'ymax':2000}
my_system.add_curve(system_parameters, name= “Curve 1”)
my_system.add_curve(system_parameters_higher_affinity, name= “Curve 2”)
my_system.show_plot()
```

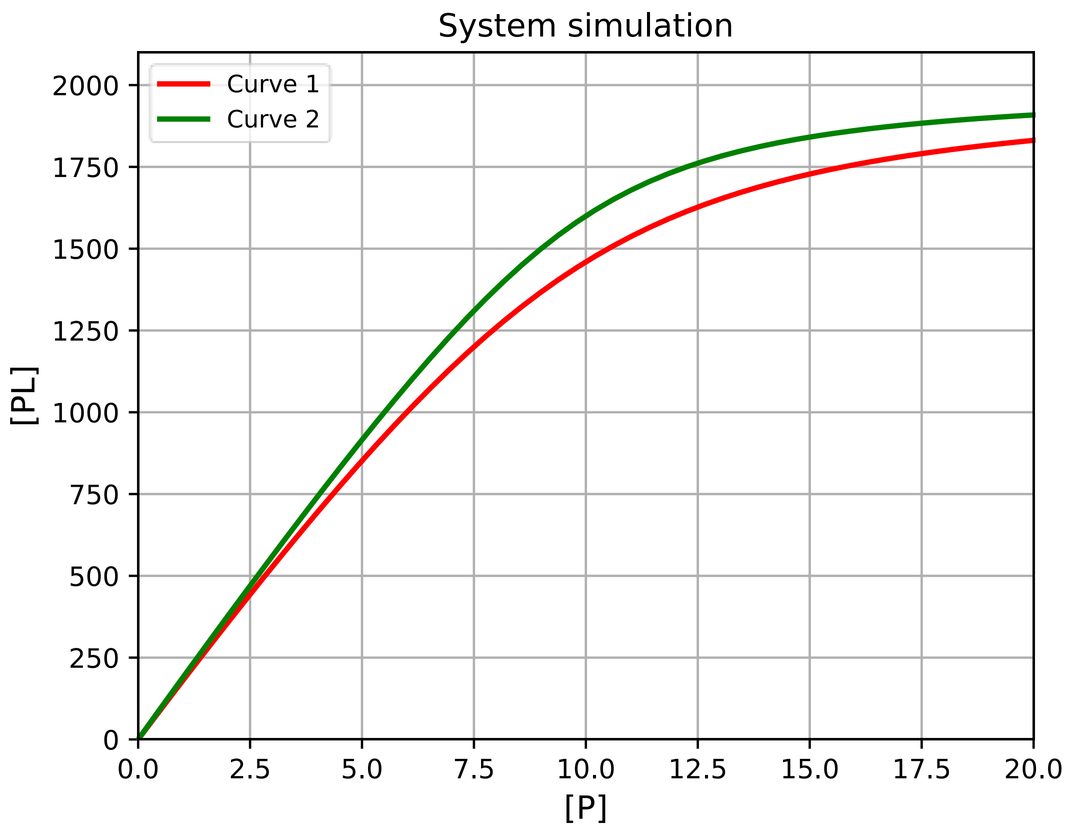

Additionally, we may transform the readout by passing a range of pbc.Readout objects to add_curve, or query. To display curves as a fraction total ligand bound, we would pass pbc.Readout.fracion_l. For more information on readouts, please see pbc.Readout in the PyBindingCurve documentation. Supplying a readout overrides the automatic signal readout selection when a ymin or ymax parameter has been found in a system:

```
system_parameters = {"p": np.linspace(0, 20), "l": 10, "kdpl": 1, 'ymax':2000}
system_parameters_higher_affinity = {"p": np.linspace(0, 20), "l": 10, "kdpl": 0.5,
'ymax':2000}

my_system.add_curve(system_parameters, readout=pbc.Readout.fraction_l)
my_system.add_curve(system_parameters_higher_affinity, readout=pbc.Readout.fraction_l)

my_system.show_plot()
```

Results in the following:

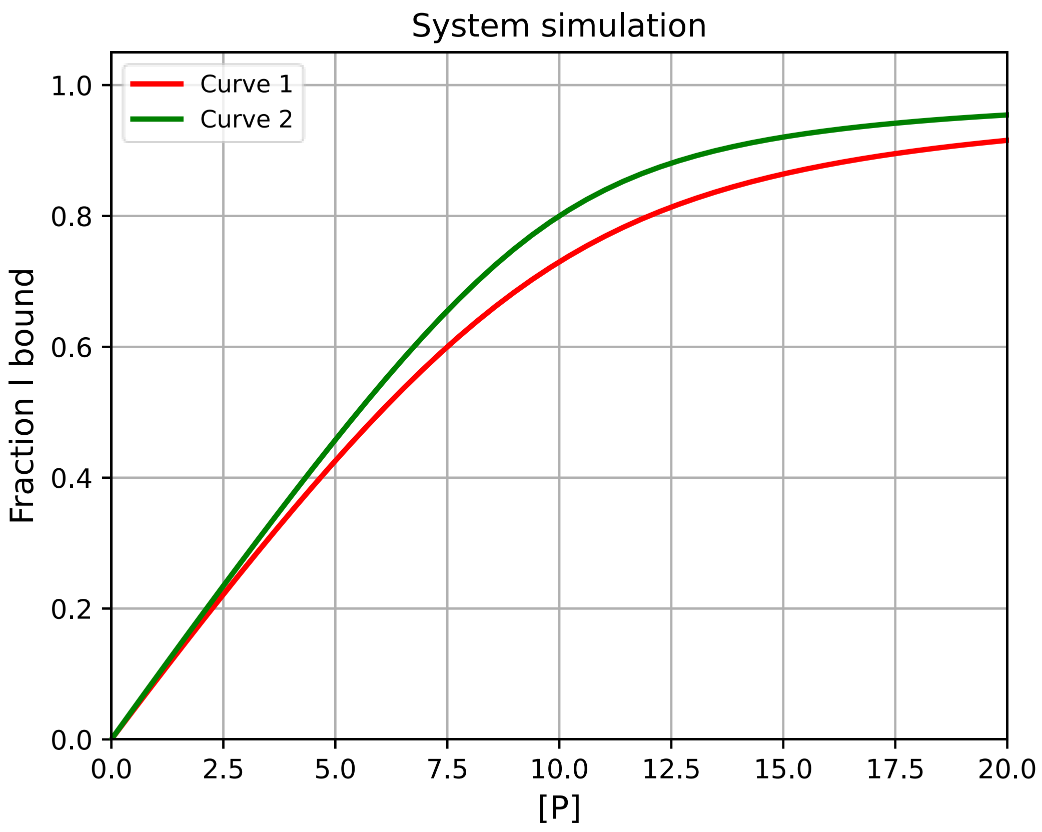
System simulation
Curve 1
Curve 2
Fraction I bound
[P]
0.0
0.2
0.4
0.6
0.8
1.0
0.0
2.5
5.0
7.5
10.0
12.5
15.0
17.5
20.0

#### Fitting of data to 1:1 binding

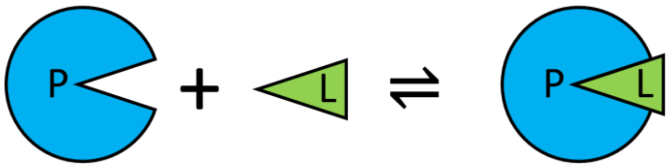

Example code is available here:

https://github.com/stevenshave/pybindingcurve/blob/master/example_1to1_fit.py

A titration of protein from nothing to 200 µM into a cuvette containing 10 µM ligand produces the following signal from the instrument:

| | | | | | | | | | | | |
|---|---|---|---|---|---|---|---|---|---|---|---|
| [P] µM | 0 | 20 | 40 | 60 | 80 | 100 | 120 | 140 | 160 | 180 | 200 |
| Signal | 54.4 | 483.2 | 636.7 | 709.3 | 798.7 | 900.5 | 907.9 | 890.6 | 901.0 | 1004.6 | 922.5 |

We may easily fit this data using PBC and obtain two values… the $K_D$ of the complex, as well as the maximal signal (ymax) achievable with the system.

We first import PBC and Numpy:

```
import numpy as np
import pybindingcurve as pbc
```

Next we create NumPy arrays to hold our protein concentration and signal:

```
xcoords = np.array([0.0, 20.0, 40.0, 60.0, 80.0, 100.0, 120.0, 140.0, 160.0, 180.0,
200.0])
ycoords = np.array([54.4, 483.2, 636.7, 709.3, 798.7, 900.5, 907.9, 890.6, 901.0, 1004.6,
922.5])
```

We now define a PBC BindingCurve object governed by the 1:1 binding system, and add the experimental data to PBC's internal plot using the add_scatter function of the returned BindingCurve object:

```
my_system = pbc.BindingCurve("1:1")
my_system.add_scatter(xcoords, ycoords)
```

When simulating a curve, we define a system with all parameters required to fully describe the simulation.  In this case, we know four things; the amount of protein in the system (xcoords), the concentration of ligand present, the minimal signal, and finally the response.  We supply the amount of protein and ligand, along with the known minimum signal for no protein being present (calculated by calling np.min on ycoords), and not yet the response to define the system which will be supplied as an argument to the fitting function in the next code segment:

```
system_parameters = {"p": xcoords, "l": 10, “ymin”:np.min(ycoords)}
```

We then perform the fit, capturing two pieces of data; the fitted system (a dictionary of system parameters best describing the system) and the fit accuracy data.  The call to PBC.BindingCurve.fit takes the known system parameters, followed by the unknown system parameters, and finally the signal data which we are fitting to.  Unknown system parameters are passed in a dictionary, much like the system parameters, but their assigned value is only used as a starting point guess for the fitting routines and can normally be set to any value. A reasonable guess at the true value, however, is good practice. Inclusion of either ymin or ymax as either a known, or unknown system parameters allows PBC to infer that we are fitting to a signal, not an absolute, known complex concentration:

```
fitted_system, fit_accuracy = my_system.fit(system_parameters, {"kdpl": 0, “ymax”:1000}, ycoords)
```

We may now print out the fitted paramers, along with the accuracy of the parameters. Internally, PBC utilises the lmfit package to return fitted parameters along with a true fit accuracy, specifying a range within which we are 95% certain that the true value is within:

```
for k, v in fit_accuracy.items():
    print(f"Fit: {k}={fitted_system[k]} +/- {v}")
```

Running the above code results in the following output:

```
Fit: kdpl=24.720148154934403 +/- 3.800620728975203
Fit: ymax=1072.308288861583 +/- 34.03937019626566
```

Indicating that the system’s KD is 24.72 +/- 3.8 µM.

To visualise how well this fit describes the experimental data, we can use the returned system parameters to plot a curve over the scatter data already added to the pbc.BindingCurve object’s internal plot. However, the returned system object currently looks like this:

```
{'p': array([  0.,  20.,  40.,  60.,  80., 100., 120., 140., 160., 180., 200.]), 'l': 10,
'ymin': 54.4, 'kdpl': 24.720148154934403, 'ymax': 1072.308288861583}
```

Simulating and plotting such a system would produce a plot with only 11 points along the x-axis and would not look correct. We therefore increase the number of points present for protein concentration with the following command:

```
 fitted_system["p"] = np.linspace(0, np.max(xcoords))
```

We may then add the curve to the plot and visualise the result.

```
my_system.add_curve(fitted_system)
my_system.show_plot()
```

Resulting in the following:

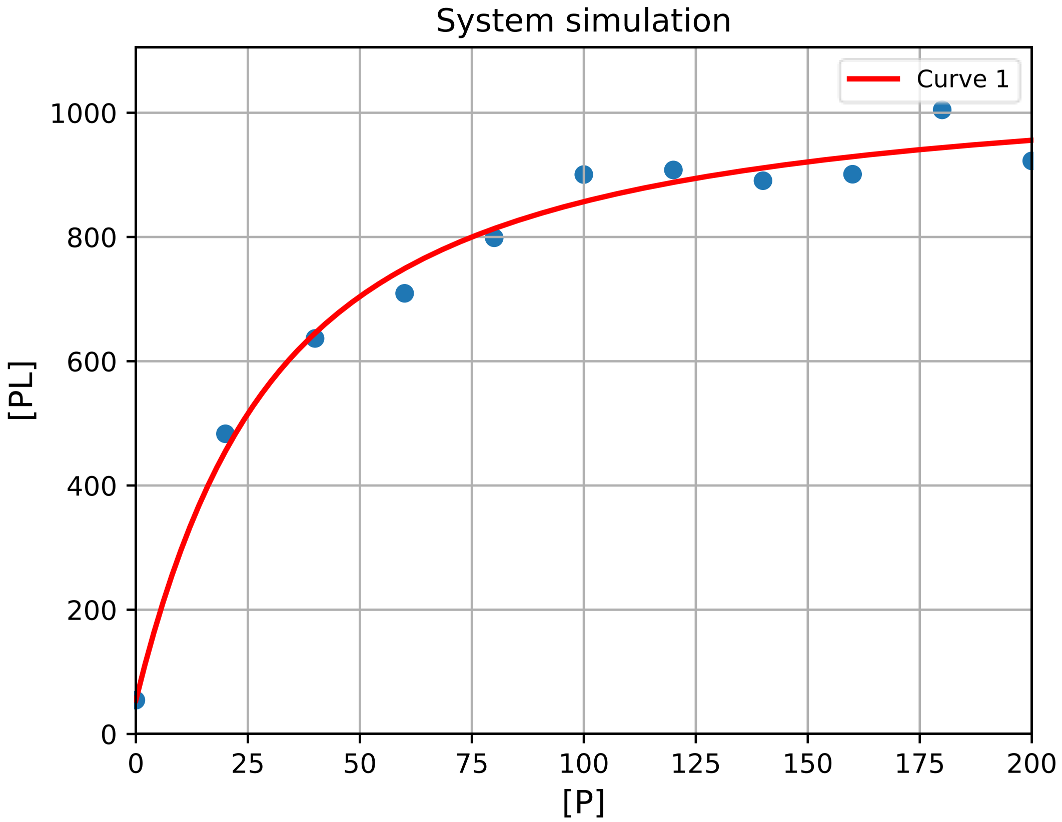
System simulation
Curve 1
[PL]
[P]
0
200
400
600
800
1000
0
25
50
75
100
125
150
175
200

#### 1:1:1 competition simulation

Competition is often used in assays, utilising displacement of a labelled ligand by new inhibitor to detect competitive binding and displacement of the label. Example code is available here: https://github.com/stevenshave/pybindingcurve/blob/master/example_competition_simulation.py

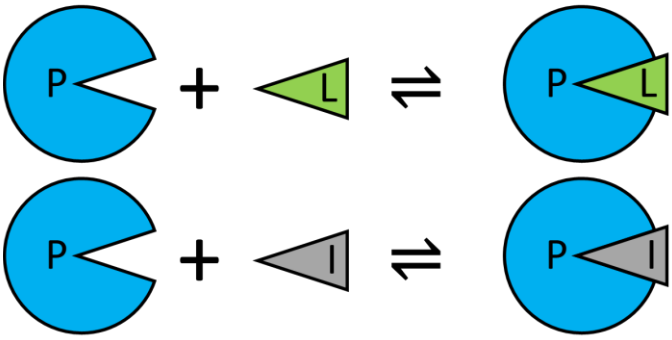

First perform imports:

```
import numpy as np
import pybindingcurve as pbc
```

We can choose to work in a common unit, typically nM, or µM, as long as all numbers are in the same unit, the result is valid. We assume µM for all concentrations bellow.

Create the PBC BindingCurve object, governed by a 'competition' system.

```
my_system = pbc.BindingCurve("competition")
```

First, let’s simulate a curve with no inhibitor present (essentially 1:1)

```
my_system.add_curve(p": np.linspace(0, 40, 20), "l": 10, "i": 0, "kdpi": 1, "kdpl": 10},
"No inhibitor")
```

Add curve with inhibitor (i)

```
my_system.add_curve(
    {"p": np.linspace(0, 40, 20), "l": 10, "i": 10, "kdpi": 1, "kdpl": 10}, "[i] = 10 µM"
)
```

Add curve with more inhibtor (i)

```
my_system.add_curve(
    {"p": np.linspace(0, 40, 20), "l": 10, "i": 25, "kdpi": 1, "kdpl": 10}, "[i] = 25 µM"
)
```

Display the plot:

```
my_system.show_plot()
```

Which results in the following:

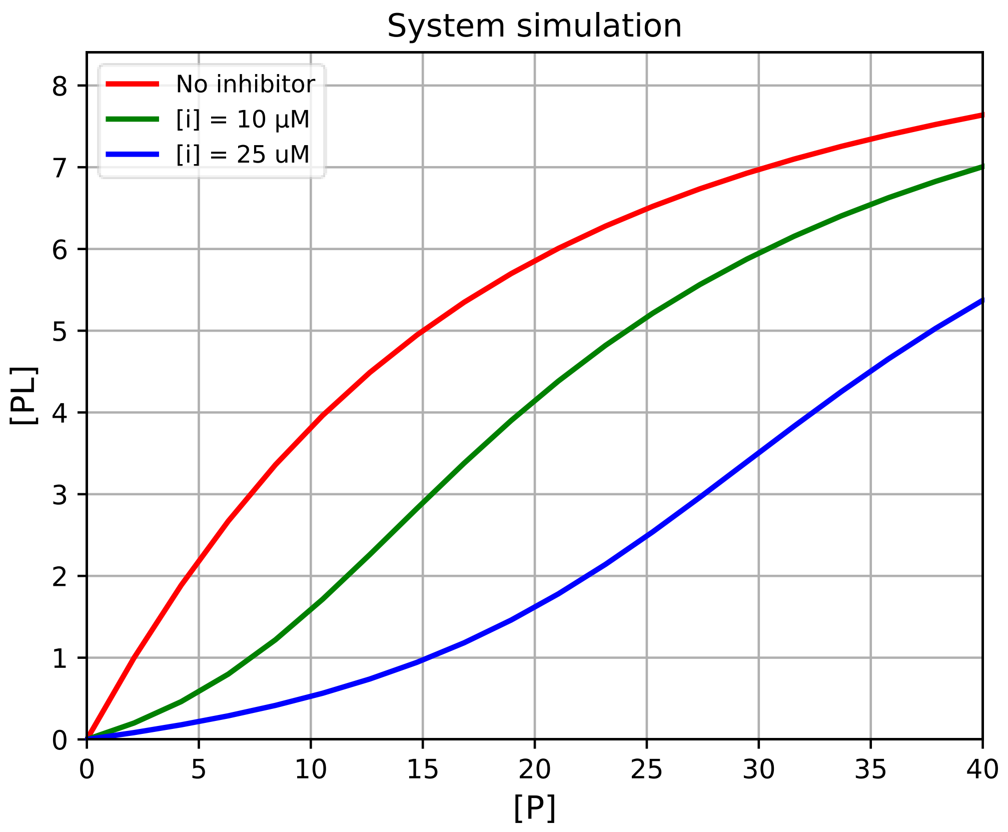
System simulation
No inhibitor
[i] = 10 µM
[i] = 25 uM
[PL]
[P]
0
1
2
3
4
5
6
7
8
0
5
10
15
20
25
30
35
40

#### Fitting to 1:1:1 competition

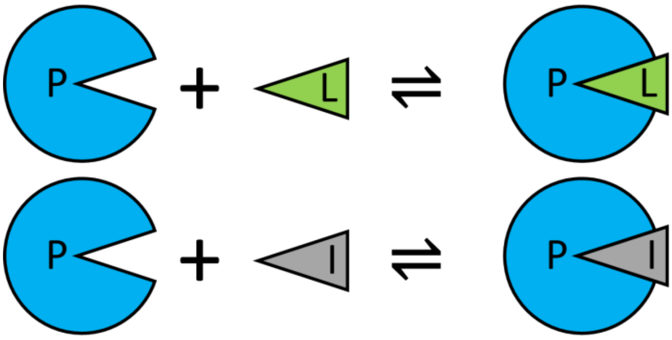

Using experimental competition data, we may obtain system parameters, such as inhibitor $K_D$. Example code is available here:

https://github.com/stevenshave/pybindingcurve/blob/master/example_competition_fit.py

Perform the standard imports:

```
import numpy as np
import pybindingcurve as pbc
```

We can choose to work in a common unit, typically nM, or µM, as long as all numbers are in the same unit, the result is valid.  We assume µM for all concentrations bellow.

Define experimental data:

```
xcoords = np.array([0.0, 4.2, 8.4, 16.8, 21.1, 31.6, 35.8, 40.0])
ycoords = np.array([150, 330, 1050, 3080, 4300, 6330, 6490, 6960])
```

Construct the PyBindingCurve object, operating on a 1:1:1 (compeittion) system and add experimental data to the plot:

```
my_system = pbc.BindingCurve("1:1:1")
my_system.add_scatter(xcoords, ycoords)
```

Known system parameters, kdpl will be added to this by fitting:

```
system_parameters = {"p": xcoords, "l": 10, "i": 10, "kdpl": 10}
```

Now we call fit, passing the known parameters, followed by a dict of parameters to be fitted along with an initial guess, pass the ycoords, and what the readout (ycoords) is:

```
fitted_system, fit_accuracy = my_system.fit(system_parameters, {"kdpi": 0}, ycoords)
```

Print out the fitted parameters:

```
for k, v in fit_accuracy.items():
    print(f"Fit: {k}={fitted_system[k]} +/- {v}")
```

Assign more points to 'p' to make a smooth plot:

```
fitted_system["p"] = np.linspace(0, np.max(xcoords))
```

Add a new curve, simulated using fitted parameters to our BindingCurve object and show the plot:

```
my_system.add_curve(fitted_system)
my_system.show_plot()
```

Which results in the following output and plot:

```
Fit: kdpi=0.44680894202996824 +/- 0.10384753604598472
Fit: ymax=9920.875421523158 +/- 98.92963212643627
```

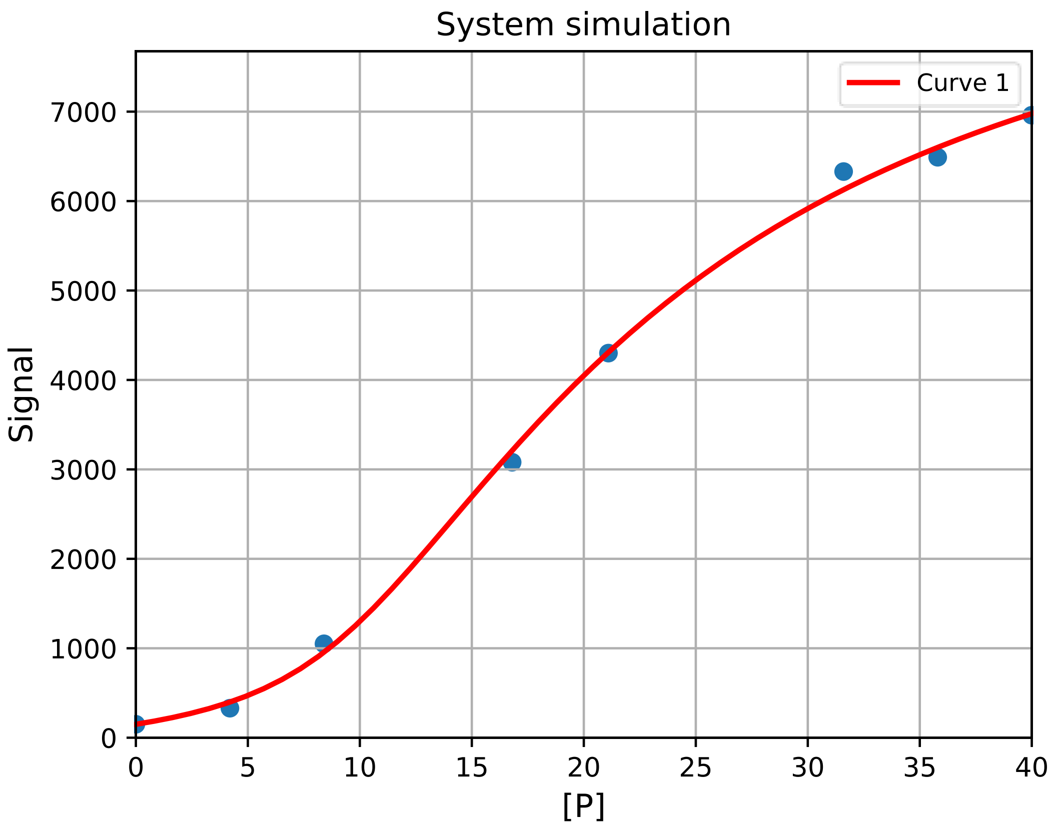

#### Homodimer formation simulation

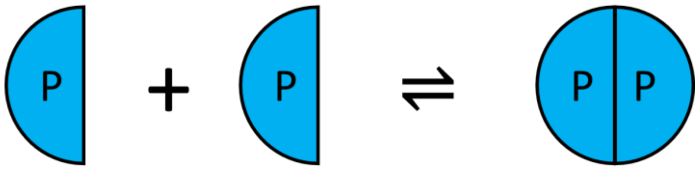

Homodimer formation is a very simple, with only one species present, two units of which bind together to make one of a new species (dimer). Dissociation of the dimer then makes two monomers. Example code is available here:
https://github.com/stevenshave/pybindingcurve/blob/master/example_homodimer_formation_simulation.py

First, we perform the standard imports:

```
import numpy as np
import pybindingcurve as pbc
```

Define our system, homodimer formation has only p (protein, or monomer concentration) and kdpp (the dissociation constant of the dimer). We can choose to work in a common unit, typically nM, or µM, as long as all numbers are in the same unit, the result is valid.  We assume µM for all concentrations bellow:

```
system_parameters = {"p": np.linspace(0, 10), "kdpp": 10}
```

Make a pbc BindingCurve defined by the 'homodimer formation' binding system:

```
my_system = pbc.BindingCurve("homodimer formation")
```

We can now add the curve to the plot and show it:

```
my_system.add_curve(system_parameters)
my_system.show_plot()
```

This produces the following simulation plot of the theoretical experiment. It is theoretical as monomer is titrated in, with no dimer present, something not achievable, although it could be done in reverse.

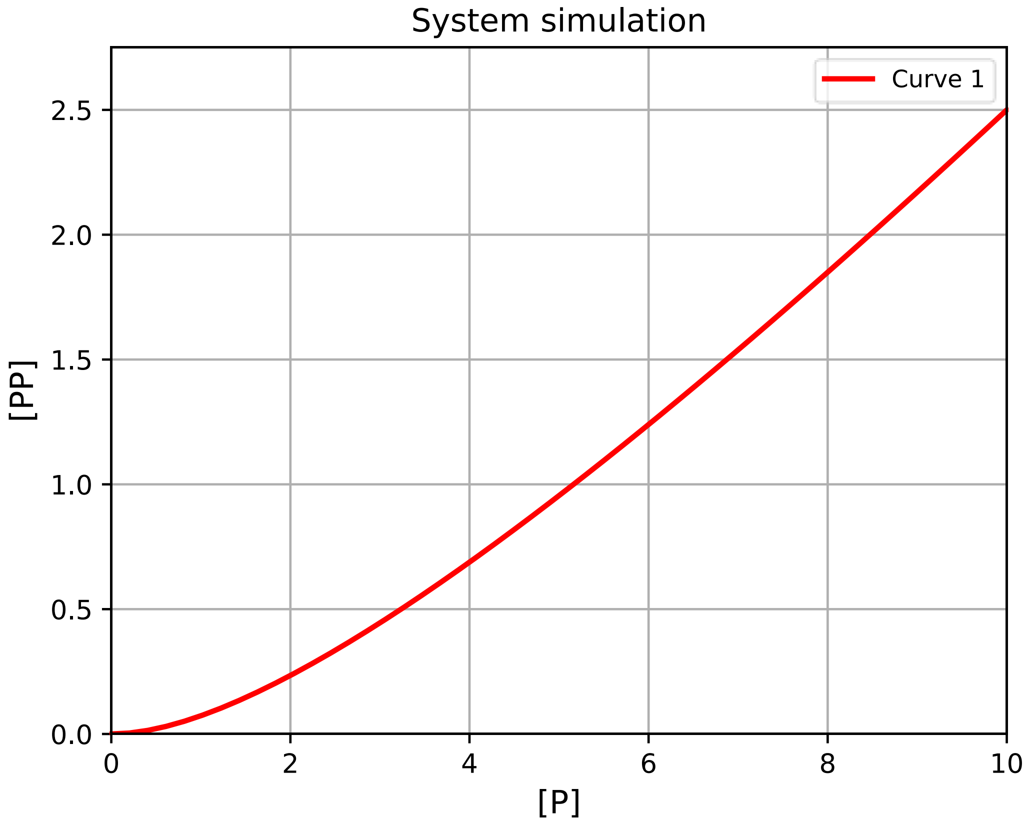
System simulation
Curve 1
[PP]
[P]
0.0
0.5
1.0
1.5
2.0
2.5
0
2
4
6
8
10

#### Fitting to homodimer formation

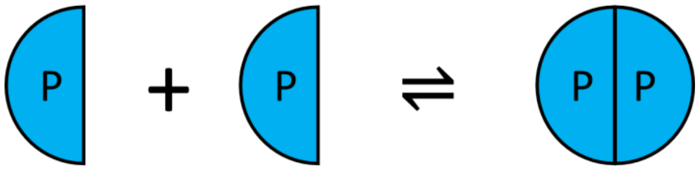

Using experimental competition data, we may obtain system parameters, such as dimerisation $K_D$. Example code is available here:
https://github.com/stevenshave/pybindingcurve/blob/master/example_homodimer_formation_fit.py

Perform the standard imports:

```
import numpy as np
import pybindingcurve as pbc
```

We can choose to work in a common unit, typically nM, or µM, as long as all numbers are in the same unit, the result is valid.  We assume µM for all concentrations bellow.

We define the known experimental data bellow:

```
xcoords = np.array([0.0,2,4,6,8,10])
ycoords = np.array([0., 0.22, 0.71, 1.24,1.88,2.48])
```

Construct the PyBindingCurve object, operating on a homodimer formation system and add experimental data to the plot:

```
my_system = pbc.BindingCurve("homodimer formation")
my_system.add_scatter(xcoords, ycoords)
```

Known system parameters, kdpp will be added to this by fitting:

```
system_parameters = {"p": xcoords}
```

Now we call fit, passing the known parameters, followed by a dict of parameters to be fitted along with an initial guess, pass the ycoords, and what the readout (ycoords) is:

```
fitted_system, fit_accuracy = my_system.fit(system_parameters, {"kdpp": 0}, ycoords)
```

Print out the fitted parameters:

```
for k, v in fit_accuracy.items():
    print(f"Fit: {k}={fitted_system[k]} +/- {v}")
```

Producing:

```
Fit: kdpp=9.939776196471206 +/- 0.15729785759220752
```

Assign more points to 'p' to make a smooth plot:

```
fitted_system["p"] = np.linspace(0, np.max(xcoords))
```

Add a new curve, simulated using fitted parameters to our BindingCurve object and show the plot:

```
my_system.add_curve(fitted_system)
my_system.show_plot()
```

Producing:

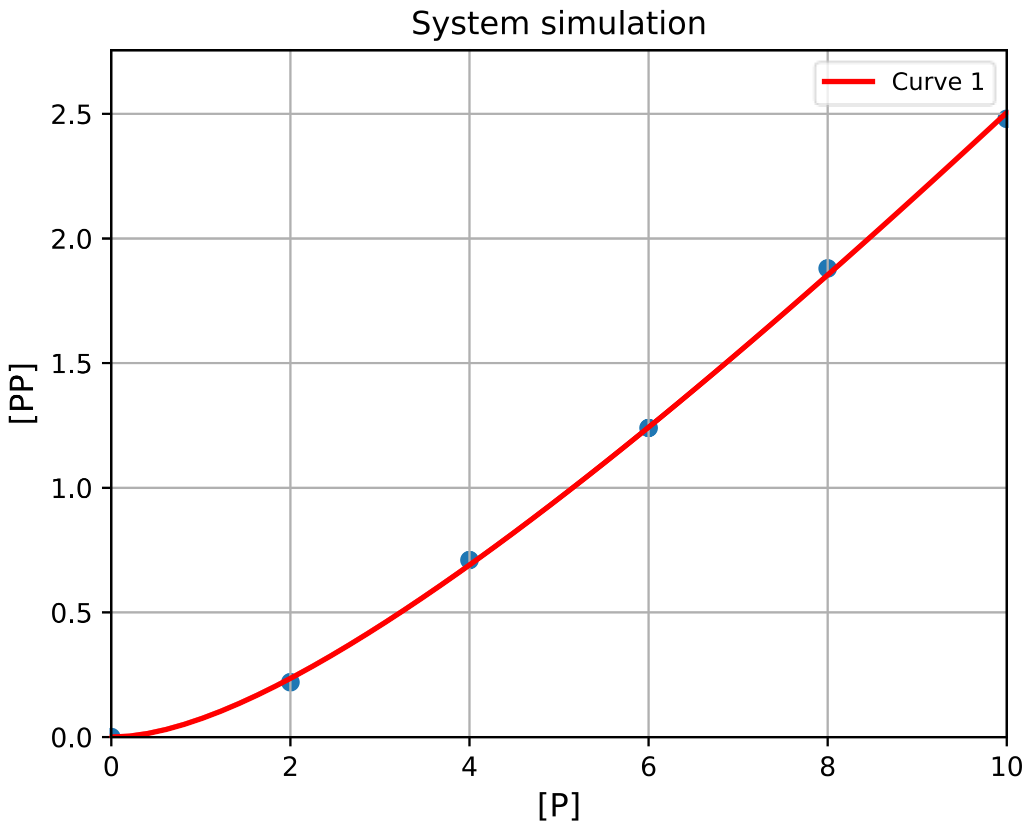

#### Homodimer breaking simulation

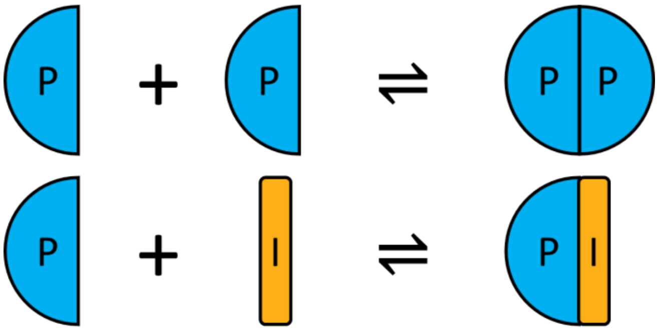

Homodimer breaking is like a competition experiment, however breaking of the dimer results in two monomers. Example code is available here: https://github.com/stevenshave/pybindingcurve/blob/master/example_homodimer_formation_simulation.py

First, we perform the standard imports:

```
import numpy as np
import pybindingcurve as pbc
```

We can choose to work in a common unit, typically nM, or µM, as long as all numbers are in the same unit, the result is valid. We assume µM for all concentrations bellow. Define out homodimer breaking system, titrating in inhibitor:

```
system_parameters = {"p": 30, "kdpp": 10, "i": np.linspace(0,60), "kdpi": 1}
```

Create the PBC BindingCurve object, expecting a 'homodimer breaking' system:

```
my_system = pbc.BindingCurve("homodimer breaking")
```

Add the system to PBC, generating a plot and display it:

```
my_system.add_curve(system_parameters)
my_system.show_plot()
```

Resulting in:

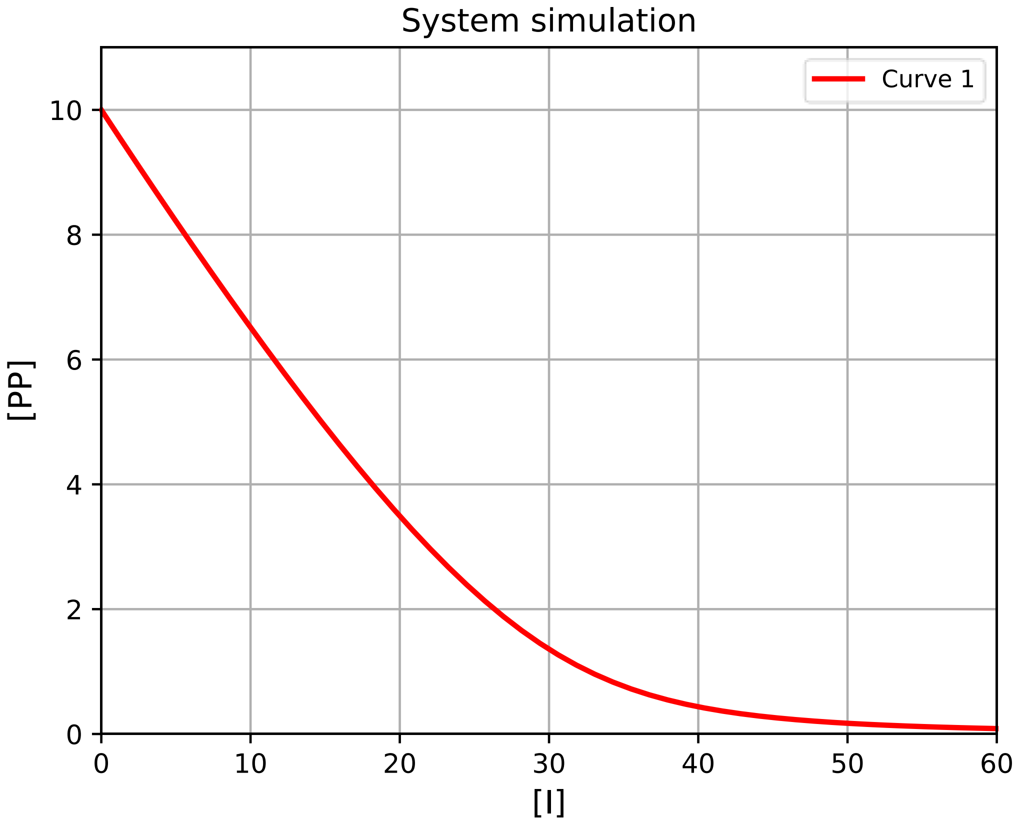
System simulation
Curve 1
[PP]
[I]
0
2
4
6
8
10
0
10
20
30
40
50
60

#### Fitting to homodimer breaking

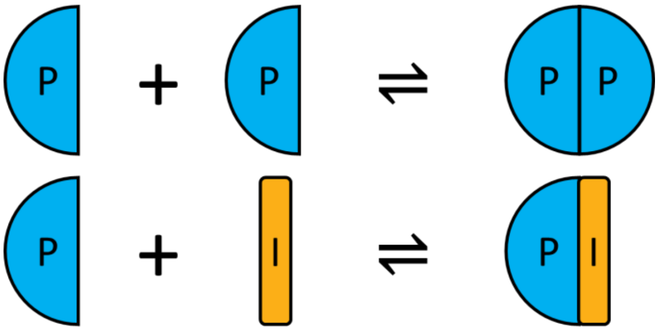

Using experimental competition data, we may obtain system parameters, such as dimerisation $K_D$. Example code is available here:
https://github.com/stevenshave/pybindingcurve/blob/master/example_homodimer_formation_fit.py

Perform the standard imports:

```
import numpy as np
import pybindingcurve as pbc
```

We can choose to work in a common unit, typically nM, or µM, as long as all numbers are in the same unit, the result is valid. We assume µM for all concentrations bellow.

Define experimental data:

```
xcoords = np.array([0.0, 16.7, 33.3, 50.0, 66.7, 83.3, 100.0])
ycoords = np.array([0.0, 0.004, 0.021, 0.094, 0.312, 1.188, 3.854])
```

Construct the PyBindingCurve object, operating on a homodimer breaking system and add experimental data to the plot:

```
my_system = pbc.BindingCurve("homodimerbreaking")
my_system.add_scatter(xcoords, ycoords)
```

Known system parameters, kdpl will be added to this by fitting:

```
system_parameters = {"p": xcoords, "i": 100, "kdpp": 10}
```

Now we call fit, passing the known parameters, followed by a dict of parameters to be fitted along with an initial guess, pass the ycoords, and what the readout (ycoords) is:

```
fitted_system, fit_accuracy = my_system.fit(system_parameters, {"kdpi": 0}, ycoords)
```

Print out the fitted parameters:

```
for k, v in fit_accuracy.items():
    print(f"Fit: {k}={fitted_system[k]} +/- {v}")
```

Assign more points to 'p' to make a smooth plot:

```
fitted_system["p"] = np.linspace(0, np.max(xcoords))
```

Add a new curve, simulated using fitted parameters to our BindingCurve object and display the plot:

```
my_system.add_curve(fitted_system)
```

```
my_system.show_plot()
```

Resulting in:

Fit: kdpi=1.0024780308947485 +/- 0.0016989355835536732

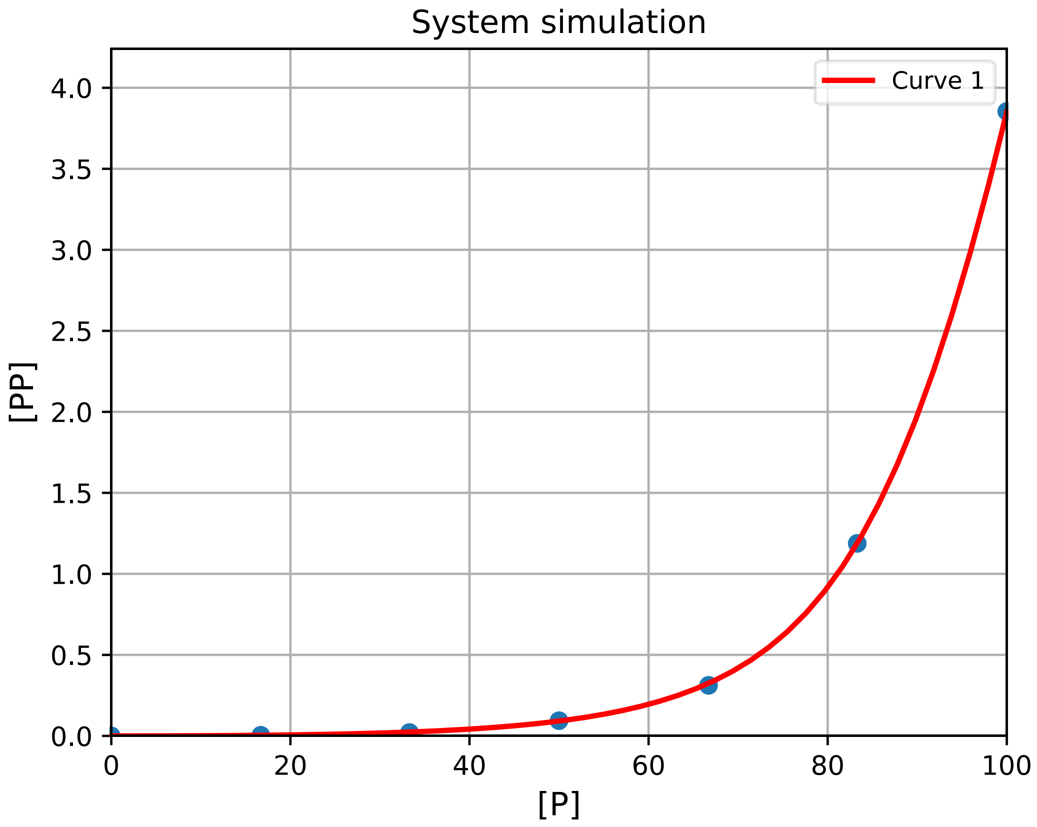

#### Simulation of custom binding systems

```
custom_system = """
    P+P<->PP*
    P+L<->PL
    PP+L<->PPL1
    PP+L<->PPL2
    PPL1+L<->PPL1L2
    PPL2+L<->PPL1L2
"""
```

Example code is available here:
https://github.com/stevenshave/pybindingcurve/blob/master/example_custom_binding_system.py

Using Lagrange multipliers as a method of solving protein-ligand binding systems, PyBindingCurve is able to write custom Lagrangian functions from very simple system definition strings. This allows the simple definition, solving, plotting and fitting to any custom system.

We define these custom systems as simple strings with reactions separated either on newlines, commas, or a combination of the two. Reactions take the form:

- `r1+r2<->p`

Denoting reactant1 + reactant2 form p.

PBC will generate and solve custom Lagrangian systems. Readouts are signified by inclusion of a star (*) on a species. If no star is found, then the first seen product is

used. Some system examples follow:

- "`P+L<->PL`" - standard protein-ligand binding

- "`P+L<->PL, P+I<->PI`" - competition binding

- "`P+P<->PP`" - dimer formation (default readout on PP - dimer)

- "`P*+P<->PP`" - dimer formation (readout specified on P - monomer)

- "`monomer+monomer<->dimer`" - dimer formation (default readout on PP)

- "`P+L<->PL1, P+L<->PL2, PL1+L<->PL1L2, PL2+L<->PL1L2`" - 1:2 site binding

KDs passed to custom systems use underscores to separate species and keep clarity in exactly what the two components are. P+L<->PL

would require the KD passed as kd_p_l. Running with incomplete system

parameters will prompt for the correct ones.

We can choose to work in a common unit, typically nM, or µM, as long as all numbers are in the same unit, the result is valid.  We assume µM for all concentrations and KDs bellow.

To simulate a highly complex system, where protein binds to ligand, but protein can dimerise, while protein dimer binds to an inhibitor, and the protein dimer has can bind a single ligand, and our readout is on the protein monomer bound to ligand, we would define the system as follows:

```
custom_system="""
       P+L<->PL*
        P+P<->PP
        PP+I<->PPI
        PPI+L<->PPIL
"""
```

We can simulate this system in Python as follows:

```
import numpy as np
import pybindingcurve as pbc
custom_system="""
        P+L<->PL*
        P+P<->PP
        PP+I<->PPI
        PPI+L<->PPIL
"""
my_system = pbc.BindingCurve(custom_system)
```

We then define the system parameters in a python dictionary.

```
system_parameters = {'p0': np.linspace(0,5),
'l0':0.5,
'i0':4.0,
'kd_p_l':1,
'kd_p_p':2,
'kd_p_p':1.2,
'kd_p_l':4.3,
'kd_pp_i':1.2,
'kd_ppi_l':0.2,
}
```

We can now add the curve to the plot. If we want multiple simulations on the same plot, then it is good to give the curve a name with the optional name parameter.

```
my_system.add_curve(system_parameters, name= “Curve 1”)
```

Finally show the plot.  Optionally, title, xlabel and ylabel variables may also be passed to title the plot and axes.

```
my_system.show_plot()
```

This produces the following plot:

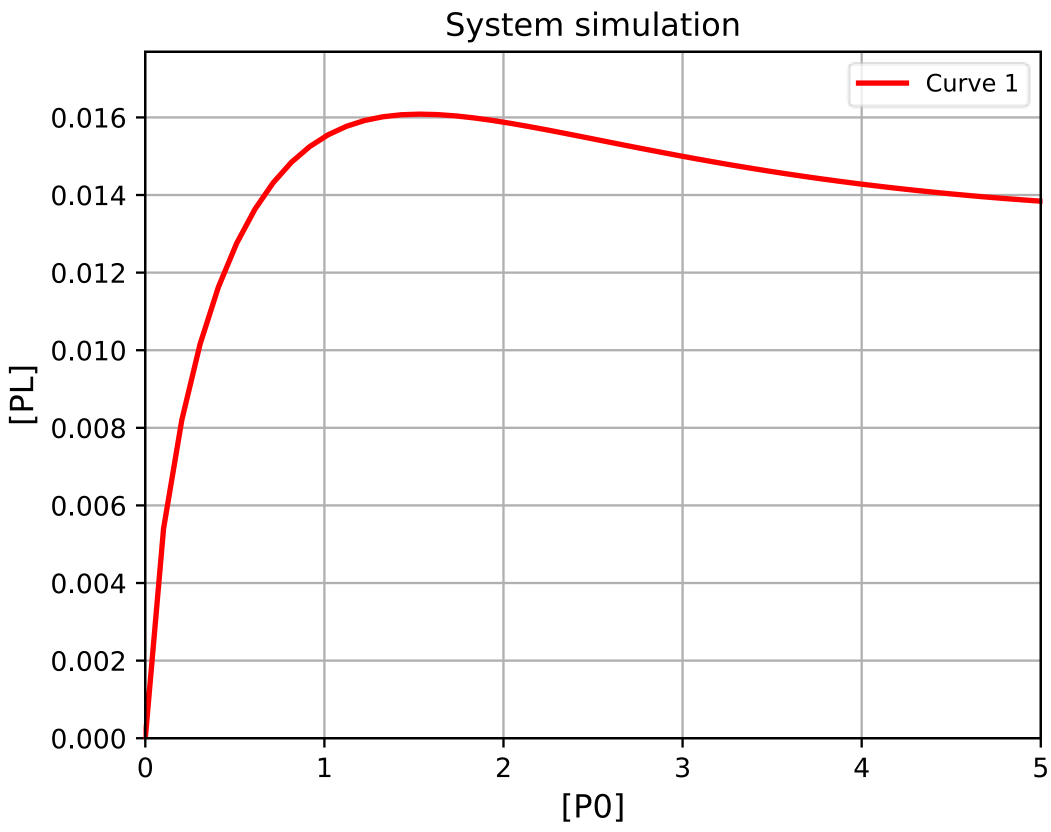

To obtain exact single points from the plot, we may call the query function of my_system:

```
print(my_system.query({'p0': 0.5,
'l0':0.5,
'i0':4.0,
'kd_p_l':1,
'kd_p_p':2,
'kd_p_p':1.2,
'kd_p_l':4.3,
'kd_pp_i':1.2,
'kd_ppi_l':0.2,
})
```

If a list or NumPy array is included as a system parameter, then a NumPy array of results is returned.

We may want to simulate a system in terms of a signal, not the concentration of complex. In this case, we may pass additional parameters, setting the ymax and/or ymin variables in the system parameters. Inclusion of these will scale the signal present between these values. This is very important if a detector is used with a maximum or minimum sensitivity and you wish to simulate response.

#### Documentation and API

Full PyBindingCurve source code can be found here: https://github.com/stevenshave/pybindingcurve

Conventionally, the standard import utilised in a run of PyBindingCurve (PBC) are defined as follows:

```
import pybindingcurve as pbc
import numpy as np
```

PyBindingCurve is imported with the short name 'pbc', and then NumPy as 'np' to enable easy specification of ranged system parameters, evenly spaced across intervals mimicking titrations.

Next, we initialise a PBC BindingCurve object. Upon initialisation, either a string or BindingSystem object is used to define the system to be simulated. Simple strings such as "1:1", "competition", "homodimer breaking" can be used as an easy way to define the type of binding system that should be mathematically modelled. See the list of available system shortcut strings bellow in the 'pbc.systems and shortcut strings' section bellow. Custom objects may also be created of type pybindingcurve.BindingSystem, of which there exist a large choice within pybindingcurve, or the user may create a custom BindingSystem to initialise PBC objects:

```
my_system=pbc.BindingCurve("1:1")
```

There are three main modes of operation within PyBindingCurve; 1) Visualisation of protein-ligand behaviour within a titration, simulating a range of conditions. 2) Simulation of a single system state with discrete parameters. 3) Fitting of experimental data. A description of these follows:

1. Simulation with visualisation: pass a dictionary containing system parameters to the add_curve function of the BindingCurve object (my_system). Required system parameters depend on the system being modelled. In the case of 1:1 binding, we require p (protein concentration), l (ligand concentration), kdpl (dissociation constant between p and l), and optionally, a ymax and/or ymin variable if dealing with simulation of a signal. add_curve expects one changing parameter, which will be the x-axis. By default, complex concentration will be the readout of a system, but that can be changed by passing different readout options to add_curve. See the 'pbc.Readout' section bellow for further information. With one curve added, we can add more curves, or simply display the plot by calling the show_plot function.

```
system_parameters={'p':np.linspace(0,20), 'l':20, 'kdpl':10)
my_system.add_curve(system_parameters)
my_system.show_plot()
```

2. Single point simulation: if you require not a simulation with a curve, but a single point with set concentrations and $K_D$s, then query may be called with a dictionary of system parameters and data returned. Additionally, if the dictionary contains a NumPy array, representing a titration (for example, the system parameters above), then a NumPy array of the readout is returned instead of a single value:

```
complex_conc = my_system.query({'p':10, 'l':20, 'kdpl':10})
```

3. Fit experimental data to a system to obtain experimental parameters, such as $K_D$. A common situation is determining KD from measurements obtained from experimental data. We can perform this as follows, with x- and y-coordinates, we add the experimental points to the plot, define system parameters that we do know (protein concentrations, and the amount of ligand),

and then call fit on the system passing in parameters to fit and an initial guess (1), along with the known parameters.  We then iterate and print the fitted parameters.

```
xcoords = np.array([0.0, 20.0, 40.0, 60.0, 80.0, 100.0, 120.0, 140.0, 160.0, 180.0, 
200.0])
ycoords = np.array([0.544, 4.832, 6.367, 7.093, 7.987, 9.005, 9.079, 8.906, 9.010, 
10.046, 9.225])
my_system.add_scatter(xcoords, ycoords)
system_parameters = {"p": xcoords, "l": 10}
fitted_system, fit_accuracy = my_system.fit(system_parameters, {"kdpl": 1}, ycoords)
for k, v in fit_accuracy.items():
    print(f"Fit: {k}={fitted_system[k]} +/- {v}")
```

#### pbc.BindingCurve

The BindingCurve object allows the user to work with a specific system, supplying tools for simulation and visualisation (plotting), querying of single point values, and fitting of experimental parameters to observation data.

##### *Initialisation*

When initialising this main class of PBC, we may supply either a pbc.BindingSystem, a human readable shortcut string such as “1:1”, “competition”, “homodimer formation”, etc, or a system definition string. For a full list of systems, shortcuts, and custom systems definition strings, please refer to the ‘pbc.systems and shortcut strings’ section.

Initialisation of a BindingCurve object takes the following arguments:

```
"""
BindingCurve class, used to simulate systems

BindingCurve objects are governed by their underlying system, defining the
(usually) protein-ligand binding system being represented. It also
provides the main interface for simulation, visualisation, querying and
the fitting of system parameters.

Parameters
----------
binding_system : BindingSystem or str
    Define the binding system which will govern this BindingCurve object.
    Can either be a BindingSystem object, a shortcut string describing a
    system (such as '1:1' or 'competition', etc), or a custom binding
    system definition string.

"""
```

Once intitialised with a pbc.BindingSystem, we may perform the following utilising its member functions.

##### *add_curve*

The add curve function is the main way of simulating a binding curve with PBC. Once a BindingCurve object is initialised,

```
add_curve(parameters, name = None, readout = None)
    """
    Add a curve to the plot

    Add a curve as specified by the system parameters to the
    pbc.BindingSystem's internal plot using the underlying binding system
    specified on intitialisation.

    Parameters
    ----------
    parameters : dict
        Parameters defining the system to be simulated
    name : str or None, optional
        Name of curve to appear in plot legends
    readout : Readout.function, optional
        Change the system readout to one described by a custom readout
        function.  Predefined standard readouts can be found in the static
        pbc.Readout class.
    """
```

##### *query*

When simulation with visualisation (plotting) is not required, we can use the query function to interrogate a system, returning either singular values, or arrays of values if one of the input parameters is an array or list.

```
query(self, parameters, readout: Readout = None):
```

```
        """
        Query a binding system

        Get the readout from from a set of system parameters

        Parameters
        ----------
        parameters : dict
            System parameters defining the system being queried.  Will usually
            contain protein, ligand etc concentrations, and KDs
        readout : func or None
            Change the readout of the system, can be None for unmodified
            (usually complex concentration), a static member function from
            the pbc.Readout class, or a custom written function following the
            the same defininition as those in pbc.Readout.

        Returns
        -------
        Single floating point, or array-like
            Response/signal of the system
        """
```

##### *fit*

With a system defined, we may fit experimental data to the system.

```
def fit(self, system_parameters: dict, to_fit: dict, ycoords: np.array, bounds: dict =
None):
        """Fit the parameters of a system to a set of data points

        Fit the system to a set of (usually) experimental datapoints.
        The fitted parameters are stored in the system_parameters dict
        which may be accessed after running this function.  It is
        possible to fit multiple parameters at once and define bounds
        for the parameters.  The function returns a dictionary of the
        accuracy of fitted parameters, which may be captured, or not.

        Parameters:
        system_parameters : dict
            Dictionary containing system parameters, will be used as arguments
            to the systems equations.
        to_fit : dict
            Dictionary containing system parameters to fit.
        xcoords : np.array
            X coordinates of data the system parameters should be fit to
        ycoords : np.array
            Y coordinates of data the system parameters should be fit to
        bounds : dict
            Dictionary of tuples, indexed by system parameters denoting the
            lower and upper bounds of a system parameter being fit, optional,
            default = None

        Returns
        -------
        tuple (dict, dict)
            Tuple containing a dictionary of best fit systems parameters,
            then a dictionary containing the accuracy for fitted variables.
        """
```

##### *add_scatter*

Experimental data can be added plots with the add_scatter command, taking a simple list of x and y coordinates

```
def add_scatter(self, xcoords, ycoords):
        """
        Add scatterpoints to a plot, useful to represent real measurement data

        X and Y coordinates may be added to the internal plot, useful when
        fitting to experimental data, and wanting to plot the true experimental
        values alongside a curve generated with fitted parameters.
```

```
        Parameters
        ----------
        xcoords : list or array-like
            x-coordinates
        ycoords : list or array-like
            y-coordinates

        """
```

*show_plot*

With curves, scatterpoints and fits applied, we may display the plot

```
    def show_plot(
            self,
            title: str = "System simulation",
            xlabel: str = None,
            ylabel: str = None,
            min_x: float = None,
            max_x: float = None,
            min_y: float = None,
            max_y: float = None,
            log_x_axis: bool = False,
            log_y_axis: bool = False,
            pbc_plot_style: dict = pbc_plot_style,
            png_filename: str = None,
            svg_filename: str = None,
            show_legend: bool = True,
        ):
            """
            Show the PyBindingCurve plot

            Function to display the internal state of the pbc BindingCurve objects
            plot.

            Parameters
            ----------
            title : str
                The title of the plot (default = "System simulation")
            xlabel: str
                X-axis label (default = None)
            ylabel : str
                Y-axis label (default = None, causing label to be "[Complex]")
            min_x : float
                X-axis minimum (default = None)
            max_x : float
                X-axis maximum (default = None)
            min_y : float
                Y-axis minimum (default = None)
            max_y : float
                Y-axis maximum (default = None)
            log_x_axis : bool
                Log scale on X-axis (default = False)
            log_y_axis : bool
                Log scale on Y-axis (default = False)
            ma_style : bool
                Apply MA styling, making plots appear like GraFit plots
            png_filename :  str
                File name/location where png will be written
            svg_filename : str
                File name/location where svg will be written

            """
```

#### pbc.systems and shortcut strings

pbc.systems contains all default systems supplied with PBC, and exports them to the PBC namespace. Systems may be passed as arguments to pbc.BindingCurve objects upon initialisation to define the underlying system governing simulation, queries, and fitting. Additionally, the following shortcut strings may be used as shortcuts:

| **Shortcut string list** | **pbc equivalent** |
|---|---|
| simple<br>1:1 | System_analytical_one_to_one__pl |
| simplelagrange<br>simple lagrange<br>1:1lagrange<br>1:1 lagrange | System_lagrange_one_to_one__pl |
| simplekinetic<br>simple kinetic<br>1:1kinetic<br>1:1 kinetic | System_kinetic_one_to_one__pl |
| homodimerformation<br>homodimer formation | System_analytical_homodimerformation__pp |
| homodimerformationlagrange<br>homodimer formation lagrange | System_lagrange_homodimerformation__pp |
| homodimerformationkinetic<br>homodimer formation kinetic | System_kinetic_homodimerformation__pp |
| competition, 1:1:1 | System_analytical_competition__pl |
| competition lagrange<br>competitionlagrange | System_lagrange_competition__pl |
| homodimerbreaking<br>homodimer breaking<br>homodimerbreakinglagrange<br>homodimer breaking lagrange | System_lagrange_homodimerbreaking__pp |
| homodimerbreakingkinetic<br>homodimer breaking kinetic | System_kinetic_homodimerbreaking__pp |
| homodimerbreakinganalytical<br>homodimer breaking analytical | System_analytical_homodimerbreaking__pp |
| 1:2<br>1:2 lagrange | System_lagrange_1_to_2__pl12 |
| 1:3<br>1:3 lagrange | System_lagrange_1_to_3__pl123 |
| 1:4<br>1:4 lagrange | System_lagrange_1_to_4__pl1234 |
| 1:5<br>1:5 lagrange | System_lagrange_1_to_5__pl12345 |

Custom systems can be passed allowing the use of custom binding systems derived from a simple syntax. This is in the form of a string with reactions separated either on newlines, commas, or a combination of the two. Reactions take the form:

- `r1+r2<->p`

Denoting reactant 1 + reactant2 form p. PBC will generate andsolve custom lagrangian systems. Readouts are signified by inclusion of a star (*) on a species. If no star is found, then the first seen product is used. Some system examples follow:

- "`P+L<->PL`" - standard protein-ligand binding
- "`P+L<->PL, P+I<->PI`" - competition binding
- "`P+P<->PP`" - dimer formation

- `"monomer+monomer<->dimer"` - dimer formation
- `"P+L<->PL1, P+L<->PL2, PL1+L<->PL1L2, PL2+L<->PL1L2"` - 1:2 site binding

KDs passed to custom systems use underscores to separate species. P+L<->PL would require the KD passed as kd_p_l. Running with incomplete system parameters will prompt for the correct ones.

#### pbc.BindingSystem

Custom binding systems may be defined through inheritance from the base class pbc.BindingSystem. This provides basic functionality through a standard interface to PBC, allowing simulation, querying and fitting. It expects the child class to provide a constructor which passes a function for querying the system and a query method. An example pbc.BindingSystem for 1:1 binding solved analytically is defined as follows:

```
class System_analytical_one_to_one_pl(BindingSystem):
    def __init__(self):
        super().__init__(
            analyticalsystems.system01_one_to_one__p_l_kd__pl, analytical=True
        )
        self.default_readout = "pl"

    def query(self, parameters: dict):
        if self._are_ymin_ymax_present(parameters):
            parameters_no_min_max = self._remove_ymin_ymax_keys_from_dict_return_new(
                parameters
            )
            value = super().query(parameters_no_min_max)
            with np.errstate(divide="ignore", invalid="ignore"):
                return (
                    parameters["ymin"]
                    + ((parameters["ymax"] - parameters["ymin"]) * value)
                    / parameters["l"]
                )
        else:
            return super().query(parameters)
```

Here, we see the parent class constructor called upon initialisation of the object with two arguments, the first is a python function which calculates the complex concentration present in a 1:1 binding system, which itself takes the appropriate parameters to calculate this. In addition, a flag is set to define when the solution is being solved analytically. The query method examines the content of the system and deals with the presence of ymin and ymax to denote a signal is being simulated. Query should ultimately end up calling query on the parent class, which has been set to return the result of the previously assigned function in the constructor.

#### pbc.Readout

The pbc.Readout class contains three static methods, not requiring object initialisation for use. These methods all take in a system parameters dictionary describing the system, and the y_values resulting from system query calls (either through simulation of querying for singular values). These readout functions offer a convenient way to transform results. For example, the readout function to transform complex concentration into fraction ligand bound is defined as follows:

```
def fraction_l(system_parameters: dict, y):
        """ Readout as fraction ligand bound """
        return "Fraction l bound", y / system_parameters["l"]
```

This returns a tuple, with the first value being used in labelling of the plot y-axis, and the second the y-values to be plotted; in this case, the original y values divided by the overall starting ligand concentration.  Similar functions can be defined and used interchangeably with those found in pbc.Readout.
